# Origin, Prospective Identification, and Function of Circulating Endothelial Colony Forming Cells in Mouse and Human

**DOI:** 10.1101/2022.07.31.502241

**Authors:** Yang Lin, Kimihiko Banno, Chang-Hyun Gil, Jered Myslinski, Takashi Hato, William C. Shelley, Hongyu Gao, Xiaoling Xuei, Yunlong Liu, David P. Basile, Momoko Yoshimoto, Nutan Prasain, Stefan P. Tarnawsky, Ralf H. Adams, Katsuhiko Naruse, Junko Yoshida, Michael P. Murphy, Kyoji Horie, Mervin C. Yoder

## Abstract

Most circulating endothelial cells are apoptotic, but rare circulating endothelial colony forming cells (C-ECFCs) with proliferative and vasculogenic activity can be cultured; the origin and naïve function of these C-ECFCs remains obscure. Herein, detailed lineage tracing reveals murine C-ECFCs emerge in the early postnatal period, display high vasculogenic potential, with enriched frequency of clonal proliferative cells compared to tissue-resident ECFCs, and are not committed to or derived from the bone marrow hematopoietic system but from tissue-resident ECFCs. In human subjects, C-ECFCs are present in the CD34^bright^ cord blood mononuclear subset, possess proliferative potential and in vivo vasculogenic function in a naïve or cultured state, and display a single cell transcriptome sharing some umbilical venous endothelial cell features like, higher Protein C Receptor and extracellular matrix gene expression. This study provides an advance for the field by identifying the origin, naïve function, and antigens to prospectively isolate C-ECFCs for translational studies.

## Introduction

Circulating endothelial cells (C-ECs) are dead or dying ECs sloughed off tissue vasculature (Blann et al., 2005; Hebbel, 2017) and their number serves as a biomarker for various disease states (Bertolini et al., 2006; Guervilly et al., 2020). In human subjects, circulating endothelial colony forming cells (C-ECFCs) with high clonal proliferative potential are a subset within C-ECs in umbilical cord or adult blood (CB) (Ingram et al., 2005; Ingram et al., 2004; Yoder et al., 2007). C-ECFCs are vasculogenic cells displaying in vivo vessel forming ability, discriminating their functions from bone marrow (BM)-derived “endothelial progenitor cells (EPCs)” (Asahara et al., 1997) restricted to playing proangiogenic roles (Kawasaki et al., 2015; Medina et al., 2017; Purhonen et al., 2008). However, several important unaddressed questions remained; where do C-ECFCs originate and do C-ECFC in their in vivo naïve state function similar to isolated, growth factor expanded, and cultured ECFC?

We have addressed these questions in mice and found murine C-ECFCs in a developmentally regulated window across several mouse strains. We identified lineage traced C-ECFCs with high proliferative potential that upon transplantation in a naive or cultured expanded state form robust vasculature. Moreover, single cell RNA sequence analysis (scRNAseq) of human umbilical cord blood (CB) distinguished various C-EC subsets. Markers defining the subsets permitted prospective isolation of C-ECFC from CB with evidence that naïve or cultured C-ECFC display clonal proliferative potential and in vivo vessel formation, defining these cells as a distinct subpopulation of C-ECs. We have thus uncovered key characteristics of C-ECFCs in mouse and man, with new information to accelerate C-ECFCs as a viable choice for cell therapy.

## Results

### C-ECFCs in Mice Possess Vasculogenic Potential Similar to Humans

C-ECFCs though present in humans, are not demonstrable in individual adult mice (Somani et al., 2007). Since CB ECFCs are enriched 60-100 fold compared to adult peripheral blood (PB) (Ingram *et al*., 2004), we isolated murine PB mononuclear cells (MNCs) from prenatal to 3 months age and seeded on OP9 feeder cells (Figure S1A), which support the ECs proliferation into spindle shaped colonies on the OP9 monolayer surface (Hirashima et al., 1999; Lin et al., 2019). Within 4 days, typical EC colonies appeared from blood of multiple strains (Figures S1B—S1G), identifying C-ECFCs in mice. The duration of C-ECFC recovery was notably short, with detectable but negligible levels prior to birth, peaking at birth, and rapidly declining in a strain-dependent manner within weeks to months (Figures 1A and S1H).

**Figure 1.**
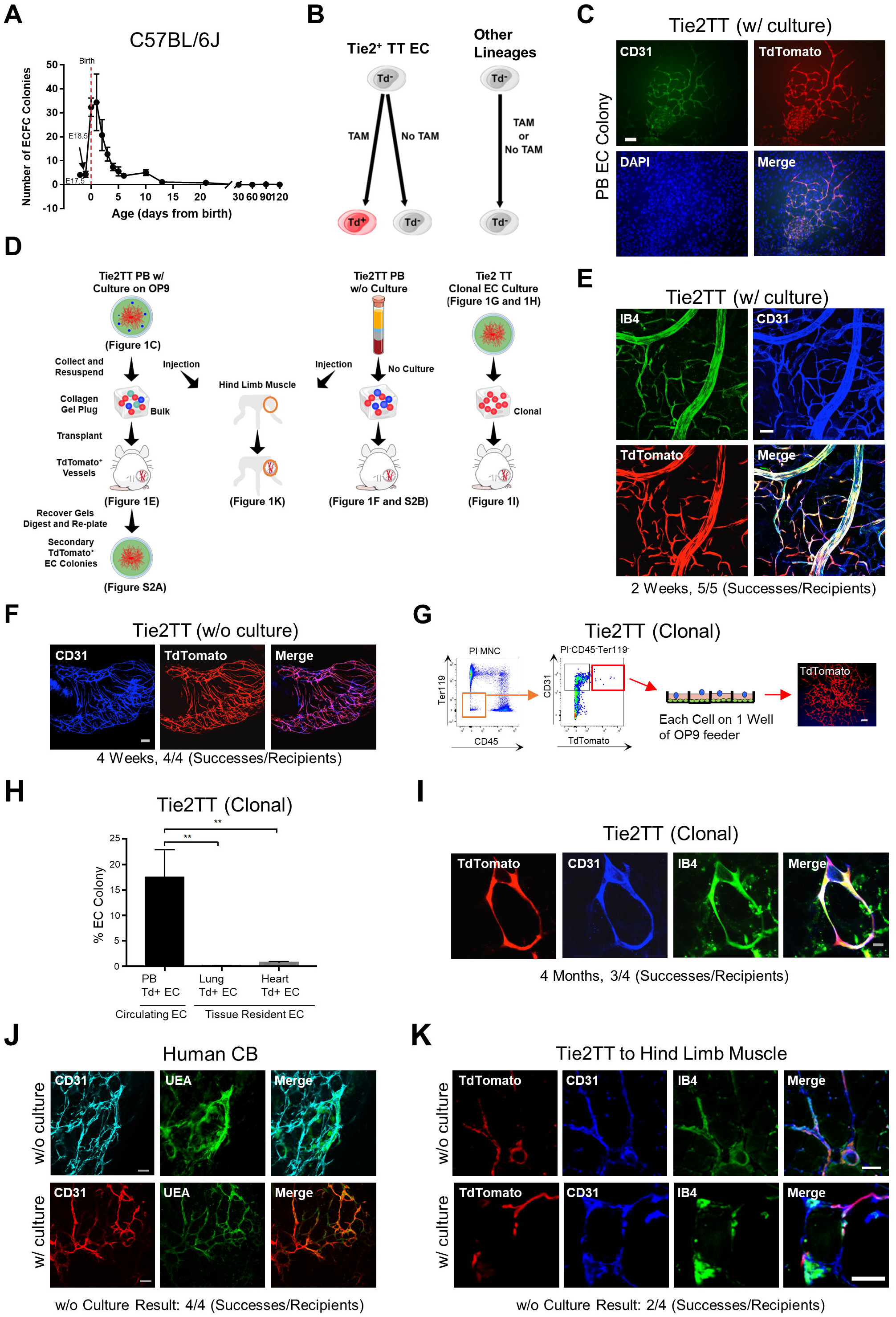
Some Murine Circulating ECFC Form Functional Blood Vessels in vivo and Have the Ability to Self-renew. (A) Kinetics of emergence of ECFC in C57BL/6J mouse blood. (B) Schematics of lineage tracing using Tie2TT mice. (C) TT^+^ EC colonies derived from PB of Tie2TT mice. (D) Schematic of collagen plug transplantation (All) and cell injection to hind limb muscle (Left and Middle) using PB-derived cells (Left), PB without OP9-culture (Middle), EC colony from clonal EC culture (Right) of Tie2TT mouse (P2). TT^+^ vessels can be digested and replated on OP9, resulting in secondary TT^+^ EC colonies (Left). (E) Two weeks after transplantation, PB-derived (with OP9-culture) TT^+^ vessels are inosculated with host vasculature (shown by systemic isolectin B4 intravenous injection; IB4). (F) Four weeks after collagen-plug transplantation using PB of Tie2TT (P2); uncultured CEC derived blood vessels (TT^+^) are shown. (G) Schematic of single cell colony forming assay using CD45^-^Ter119^-^CD31^+^ TT^+^ cells in Tie2TT mice (P2). Right panel shows a representative picture of an endothelial colony. (H) Quantitation of the frequency of ECFCs from PB, lung, or heart derived CD45^-^Ter119^-^CD31^+^ TT^+^ cells in Tie2CreTT mice (P2). (I) “Single Tie2CreTT TT^+^ cell derived EC colony”-derived blood vessel in vivo 4 months after transplantation. (J) Uncultured and cultured human CB CD34^+^CD45^-^ cells (MACS-sorted) can form functional blood vessels in vivo after transplantation. (K) Representative TT^+^ vessel of uncultured and cultured Tie2TT PB injection to hind limb muscle. Scale bars represents 200 µm in (C), (E), (F), (G), (I), and (J). 50 µm in (K). **, p<0.01

To trace C-ECFCs, we generated a Tie2CreERT2:Rosa-TdTomato (Tie2TT) mouse, which conditionally expresses TdTomato (TT) under control of Tie2 promoter confirmed to be endothelial lineage restricted (Hochstetler et al., 2019) (Figure 1B). Neonatal mice (P0−P3) were treated with Tamoxifen (TAM) and P4 PBMNCs were plated on OP9 cells. TT^+^ EC colonies were efficiently formed in 155/177 (88%) of wells (Figures 1C and 2A). Recovered PB-derived TT^+^ EC colonies with rare surviving TT^−^ HC were suspended in a collagen gel plug and implanted subcutaneously in NOD/SCID mice. Only TT^+^ vessel formation was observed in all gels after 2 weeks (Figures 1D and 1E). Importantly, the donor TT^+^ vessels inosculated with host vessels and were perfused with the IB4 lectin intravenously infused (Figure 1E) within 2 weeks of implantation. After cells from vessels were recovered with enzymatic digestion and replated on fresh OP9 cells, secondary TT^+^ EC colonies were identified from every plug tested (Figure S2A). Thus, most C-ECFCs can be labeled with the Tie2TT tracing system and possess prominent clonal proliferation and functional in vivo vessel forming potential.

**Figure 2.**
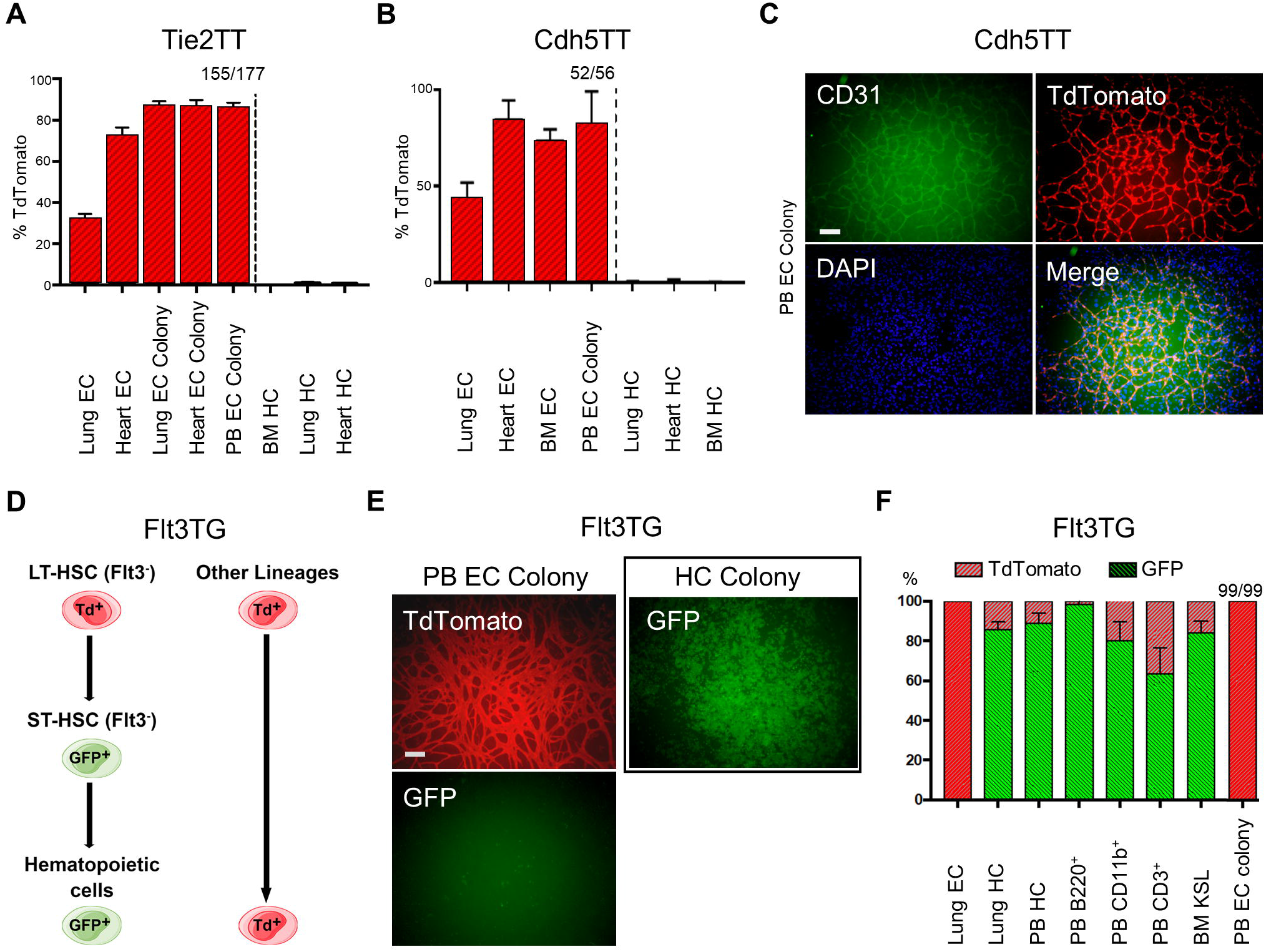
Murine C-ECFCs Are Derived from Resident ECs. (A and B) Quantitation of labeling efficiency of endothelial and hematopoietic lineage, and resultant EC colonies in each tissue/organ from Tie2TT (A) and Cdh5TT (B) mice. (C) TT^+^ EC colonies derived from PB of Cdh5TT mice. (D) Schematics of lineage tracing using Flt3TG mice. (E) EC colonies (left) and HC colonies (right) derived from PB of Flt3TG mice. (F) Percentage of GFP and TT expressing cells in the fraction of lung EC, lung HC, BM KSL, PB HC, PB B220^+^, PB CD11b^+^, and PB CD3^+^. All EC colonies derived from peripheral MNCs are TT^+^. Scale bars represents 200 µm. KSL, c-Kit^+^Sca1^+^Lineage-cell.

### Murine and Human Naïve C-ECFCs Form Functional Blood Vessels and Display Ability to Self-renew

C-ECFCs’ vasculogenic potential was subsequently tested without any in vitro cell culture step to address any criticism that cultured ECFCs are an artifact of tissue culture adaptation (Figure 1D). Freshly isolated Tie2TT PBMNCs formed TT^+^ vessels when implanted in mice (Figures 1F and S2B) reflecting the inherent (non-induced) vasculogenic nature of naïve C-ECFCs present in circulating murine blood. Next, to assess the clonal proliferative properties, Tie2TT^+^ cells (TAM at P0−P1) in PBMNCs, lung, and heart ECs were plated at a single cell level (at P2) (Figures 1D and 1G). EC colonies formed in 17% of wells from PBMNCs, more than 20 times greater frequency than lung or heart ECs (Figures 1H and S2C−S2E), suggesting postnatal blood is enriched with C-ECFCs. However, tissue Tie2TT^+^ vessel derived ECFCs possess clones with proliferative potential that is greater (>10,000 cells/colony) than C-ECFC, where the most proliferative clones produced 2001-10,000 cells (Figures S2C and S2E). These results suggest C-ECFC are derivatives of tissue ECFC since the known proliferative hierarchy of ECFC dictates that the most proliferative ECFC give rise to clones restricted to fewer cells/colony (Figure S2E) (Ingram *et al*., 2004; Mund et al., 2012).

EC colonies formed from single plated TT^+^ ECs were recovered and implanted, and TT^+^ vessels were identified and survived for up to 4 months (Figure 1I). This indicates some single Tie2^+^TT C-ECFC display remarkable proliferative and in vivo vessel forming potential and evidence that C-ECFC are derivatives of resident tissue Tie2^+^TT ECFC. Similar to C-ECFCs in postnatal mice, human CB ECFCs form a hierarchy of clonal EC colonies and display vasculogenesis in vivo (Ingram *et al*., 2004; Mund *et al*., 2012). We now report freshly isolated CD34^+^CD45^−^ CBMNCs exhibit vasculogenesis in all gels after transplantation without prior in vitro culture comparable to CB ECFC isolated and expanded in culture (Figure 1J). Moreover, cells recovered from the formed vessels give rise to secondary ECFC colonies upon replating (Figure S2F). Thus, circulating CB ECFC form vessels in vivo, which by nature holds endothelial intima in quiescence by endocrine, autocrine, paracrine, and mechanical inputs (Ricard et al., 2021), but within which self-renewing ECFC were maintained that produced secondary ECFC upon plating in vitro.

To demonstrate whether naïve murine C-ECFCs contribute to tissue vasculature in murine tissue, we injected PB (pooled blood of 6 neonates, P2) of Tie2TT mice (TAM at P0−P1) directly into the hind limb muscle of host C57Bl/6J mice. TT^+^ EC engraftment was confirmed in the tissue injected with PB itself (2/4 recipient mice) including evidence that the vessels were functionally active, inosculated to host vessels, and perfused with systemically injected IB4 lectin (Figure 1K). These findings highlight naïve C-ECFCs directly contribute to vascular regeneration in murine tissues in vivo.

### Murine C-ECFCs Are Derived from Resident ECs

While TT specifically labeled ECs, but not HCs, in organs (Figure 2A), we asked whether other vascular endothelial or blood cell restricted reporters marked C-ECFC. Cdh5TT mice conditionally express TT under control of the endothelial specific Cdh5 promoter (Wang et al., 2010) and most tissue-resident ECs were TT^+^ but HCs were TT^-^ (Figure 2B). Culturing PB from Cdh5TT mice on OP9 cells led to identification of TT^+^ EC colonies in 52/56 (93%) of wells (Figures 2B and 2C), suggesting C-ECFCs are derived from vascular ECFCs that constitutively express Tie2 and Cdh5. Indeed, resident vascular endothelial stem/progenitors (VESP) with ECFC potential are known to reside within the vascular endothelial intima and have been shown to play a major role in vascular regeneration (McDonald et al., 2018; Wakabayashi et al., 2018).

Next, we confirmed the endothelial origin for C-ECFC and not a BM hematopoietic origin (as claimed for EPC (Asahara *et al*., 1997)) using the hematopoietic-restricted Flt3cre; mTmG reporter mouse (Boyer et al., 2011) (Figures 2D and S2G). In this animal, TT is ubiquitously expressed in all cells, except those cells that express Flt3cre, since Cre-mediated recombination irreversibly switches TT to GFP fluorescence. As a result, Flt3^+^ short-term hematopoietic stem cells (HSC) and all of their hematopoietic progeny express GFP (Figures 2D, 2F and S2H) (Tarnawsky et al., 2018). As anticipated, HC colonies from Flt3cre; mTmG PBMNC were labeled with GFP (Figure 2E). The MNCs isolated from 25 individual Flt3cre; mTmG mice formed a total of 99 EC colonies (PB EC colony) and not one expressed GFP, but all colonies were TT^+^ (Figures 2E and 2F), similar to control primary lung EC. Thus, C-ECFCs are not committed to or derived from the BM hematopoietic lineage but are marked by typical endothelial transgenic markers.

### Single Cell RNA Sequencing Reveals Unique Endothelial Cell Related Clusters in Circulating Human Cord Blood Mononuclear Cells

To discriminate human C-ECFCs from other blood cells, we compared scRNAseq of fresh CBMNC and HUVEC isolated from matching umbilical cords. (Figure 3A and 3B). We annotated hematopoietic lineage clusters (Group 1-1 and 1-2, Myeloid; C10, NK and T; C12, Mk; C13, B; C15, HSPC) based on characteristic genes (Figures 3A, 3C and S3A) (Novershtern et al., 2011; Zhao et al., 2019). A second major cell population (group 2) consisted of 6 clusters (C3, 4, 5, 7, 14, 16) expressing EC markers such as PECAM1, CDH5, and VWF, but not HC genes (Figures 3B−3D). SCENIC analysis (Aibar et al., 2017) identified FOXC2, FOXC1, and NR2F2 transcriptional factor (TF) regulons as significantly enriched in C16 (Figure 3E and Table S1). C3-5, and 7 showed significant enrichment of KLF2, KLF4, and NFKB2 regulons, indicative of response to blood shear stress (Figure 3E) (Feaver et al., 2013). In C14, most EC-related TF regulons were repressed (Table S1), and higher globin and glycophorin expression were observed (Figure S3B), suggesting C14 may represent rare hemangioblastic EC present during pregnancy (Aplin et al., 2015). Such results suggest C16 as a candidate of C-ECFCs; an early differentiated cluster with enriched expression of TFs of endothelial development, moderate expression of typical EC markers, and increased Protein C receptor (PROCR) expression (Figure 3D). Volcano plots depict the enrichment of ALDH1A1 and ECM genes, such as EFEMP1, BGN, and DCN (Figure 3F). To further investigate the relationship between C16 and HUVEC, we integrated CBMNC and HUVEC UMAPs (Figure 3G). C16 were located closest to or inside the large cluster groups created by HUVECs (41/45 cells) with the majority of C16 forming a cluster with HUVECs (25/41 cells) (Figure 3G). Notably, while ECM genes and PROCR were expressed highly in HUVECs, only C16 among CBMNCs retained their expression, albeit at a reduced level (Figure 3H). Finally, the trajectory analysis of integrated CBMNC ECs and HUVECs showed HUVECs were present at earlier times than most CBMNC ECs, and C16 was clearly shown to be in the transitional path between the two (Figure 3I). Taken together, these results suggest C16 (C-ECFC candidate) display transcriptome features resembling HUVECs, more than any other CBMNC clusters, and trajectory evidence for derivation from resident vascular endothelial cells.

**Figure 3.**
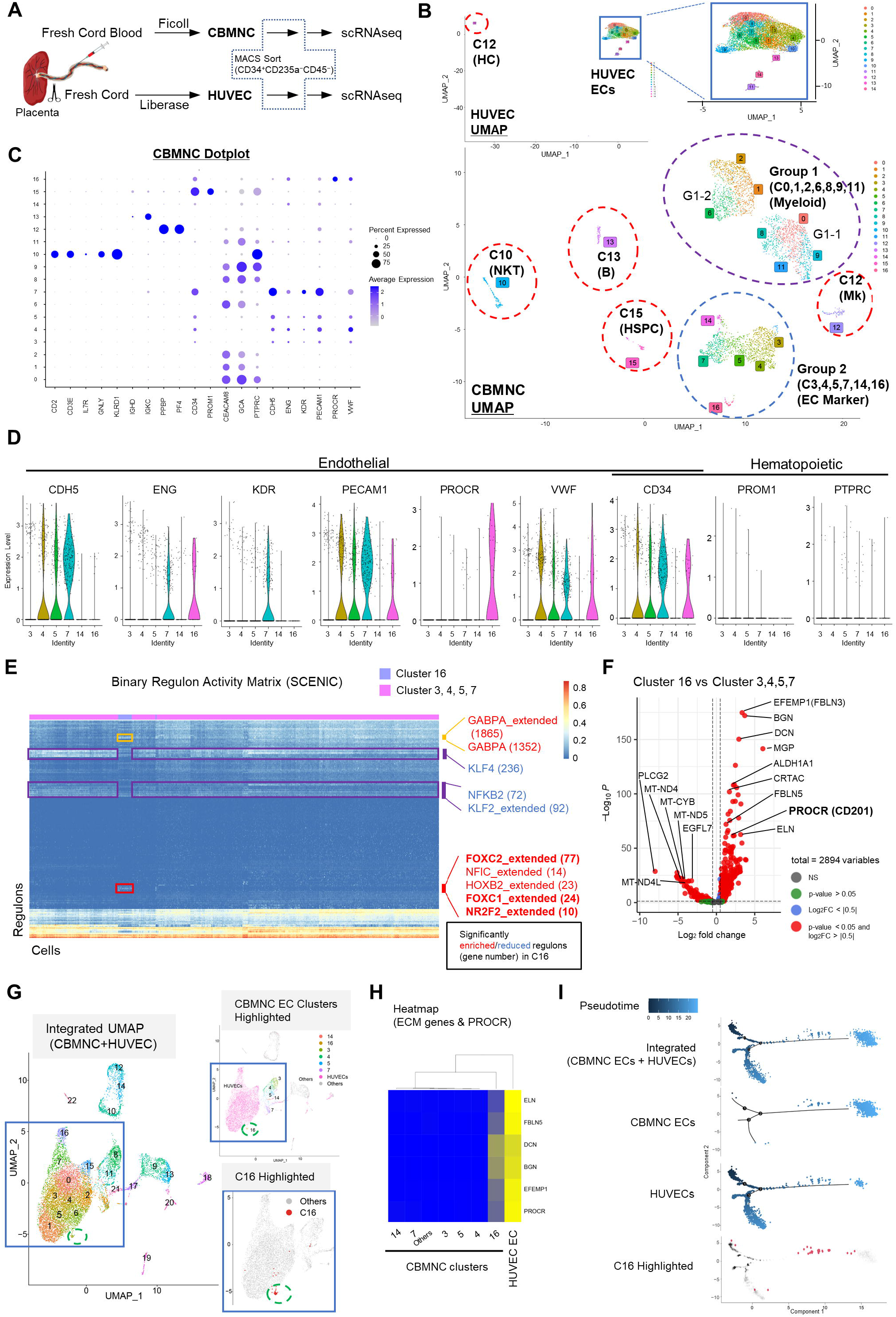
Single Cell RNA sequencing Reveals Unique Endothelial Cell Related Clusters in Circulating Cord Blood Mononuclear Cells. (A) Schematic of scRNAseq assay using freshly isolated CBMNC and HUVEC. CD34^+^ CD235a^−^CD45^−^ cells are enriched by MACS. (B and C) Global UMAP plots of CBMNCs and HUVECs (B), and dot plots of scaled average expression of major canonical markers (columns) in all clusters of CBMNCs (rows) (C). (D) Violin plots of EC genes comparing EC Marker-expressing clusters of CBMNC. (E) SCENIC analysis result on CBMNC clusters. Binary regulon activity matrix is shown with AUCell scores, indicating the activity of each TF regulon in each cell. Some of significantly enriched or reduced TF regulons in C16 are shown in right (with related gene number). (F) Volcano plots comparing C16 vs C3,4,5,and 7. Representative genes are shown. PROCR is the only cell surface marker gene in the plot with significant p-value. (G) Integrated UMAP of CBMNC and HUVEC (left). CBMNC EC clusters and C16 highlighted (middle and right), respectively. (H) Heatmap of mean expression of ECM genes and PROCR in annotated clusters. (I) Pseudotime analysis using CBMNC EC (C3-5, 7, 14, 16) and HUVEC (w/o C12), showing integrated plot and sample plot. C16 (red) is highlighted in below. See also Figure S3 and Figure S4.

### Colony-Forming Potential of PROCR-high and CD34-bright Endothelial Population in CBMNC Implicates the Existence of Circulating Endothelial Colony Forming Cells

Confirmation of scRNAseq results using colony assays demonstrated appearance of EC colonies only from PROCR enriched fraction (Figures 4B, S4A and Table S2), indicating PROCR is a novel marker for C-ECFC. The capacity of CD34^+^ to identify C-ECFCs has been reported by several groups (Mund *et al*., 2012; Tura et al., 2013), and herein, we have demonstrated uncultured naïve CD34^+^ populations form vessels, like cultured cells, in vivo (Figure 1J). MACS selected CD34^+^ fractions identified “CD34^bright^ cells” in all CB samples (n=17) (Figure S4B). Recently, C-ECs have been reported to be present in the CD34^bright^ population, not in CD34^high^ or CD133^+^ population (Case et al., 2007; Estes et al., 2010; Lanuti et al., 2016). Here, we discovered a method to magnetically enrich the CD34^bright^ population (Figure 4A). The highest CD34 expressing cells failed to be released from the magnetic beads after incubating the cells for the manufacturers instructed time and we discovered that C-ECFCs only emerged from this CD34 beads-unreleased fraction; the CD34^bright^ enriched population (Figure 4C and Table S2).

**Figure 4.**
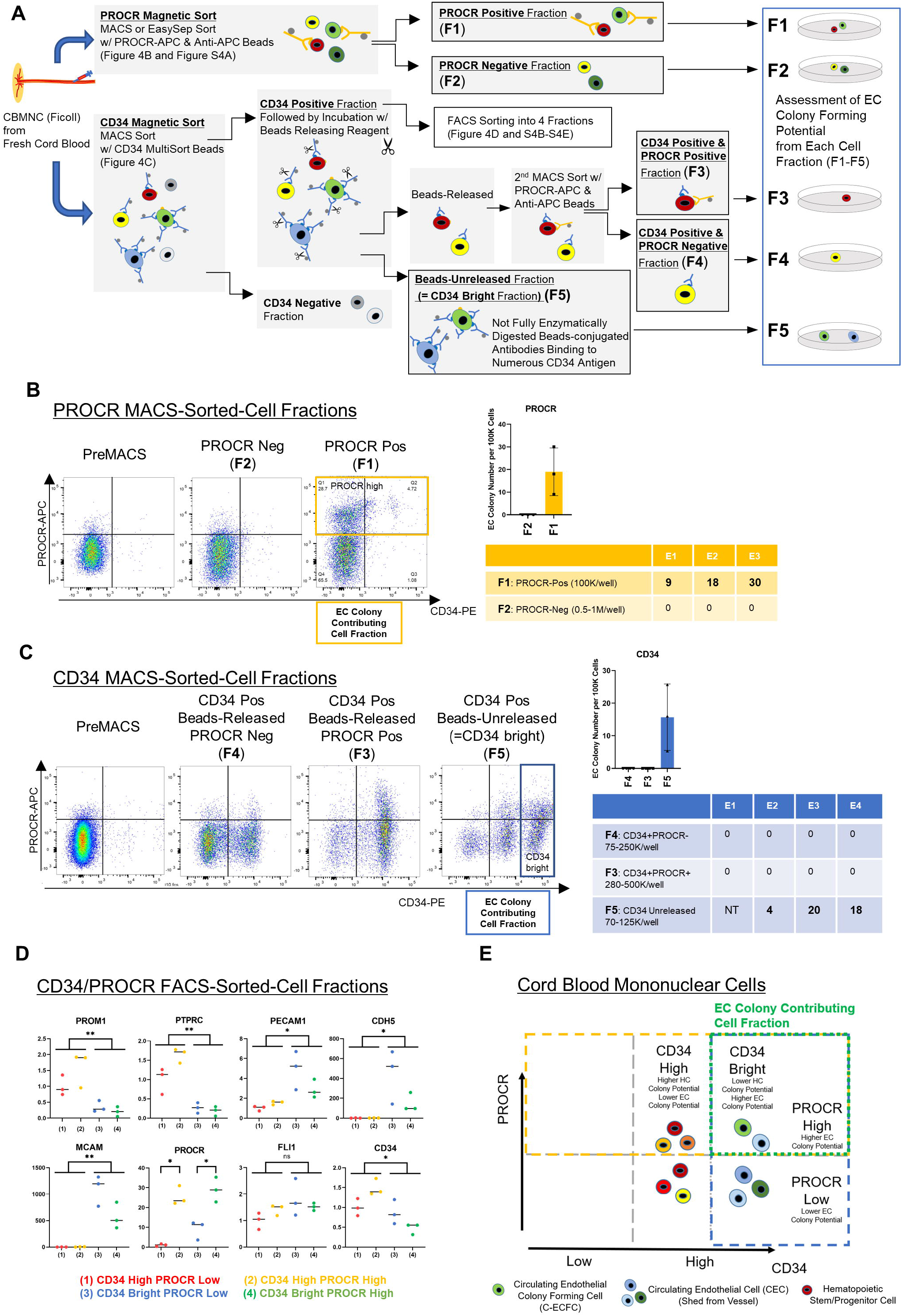
Colony-Forming Potential of PROCR-high and CD34-bright Endothelial Population in CBMNC Identifies C-ECFCs within CECs. (A) Schematic of in vitro EC colony-forming assay using fresh CB. After Ficoll separation, Above: cells are magnetic sorted with PROCR-APC & Anti-APC beads, followed by culture using cells from each fraction (F)1 and F2. Below: Cells are magnetic sorted with CD34 MultiSort Beads and CD34-Positive fraction is incubated with beads-releasing reagent. Beads-released CD34 positive fraction is further sorted with PROCR-APC and Anti-APC beads, leading to F3 and F4, and followed by culture. Beads-unreleased CD34 positive fraction (enriched with CD34^bright^ population) (F5) is also cultured. (B, C) Flow cytometry analysis of pre- and post-MACS for each fraction (F1-F2 in B, F3-F5 in C). Quantitation of the number of EC colonies per 100K cells (right graph). Table shows seeding cell fraction (with cell number per well) and EC colony number in each experiment (E). NT; Not tested. (D) Fold changes of the expression of EC and HC related genes in CD34 bright/high with PROCR high/low fractions after FACS sorting. (E) Schematic of presumed roles from each fraction using CD34 and PROCR expression in flow cytometry based on the results of the EC colony contributing cell fraction from Figure 4B (yellow dots) and Figure 4C (blue dots). Error bars represent SD. Student’s t tests were used to compare two groups. ns, not significant; *, p<0.05; **, p<0.01. See also Figure S4.

Since PROCR expression presents in CD34^high^ and CD34^bright^ subsets, functional properties of the four fractions were tested following FACS isolation (Figure S4B). A smaller subset of the CD34^bright^ population formed HC colonies (Figure S4C) compared to CD34^high^ fraction. On the contrary, EC colonies were observed only in the CD34^bright^PROCR^high^ fraction (1 out of 3) and otherwise no colonies were observed (Figures S4D and S4E), consistent with the known blunting of ECFC colony formation encountered by FACS sorting (Mund *et al*., 2012). As supporting evidence, quantitative gene expression analysis revealed significantly lower PROM1 and PTPRC, and higher EC marker expression in CD34^bright^ cells. CDH5, PECAM1, and MCAM expression was lower in the PROCR^high^ fraction, suggesting this fraction may be more immature (Figure 4D). In sum, our studies showed the CD34^bright^ population is representative of C-ECs, of which the PROCR^high^ subset displays features of C-ECFCs that possess clonal proliferative potential (Figure 4E).

## Discussion

We report that murine C-ECFC display evidence of emergence from tissue resident vessel ECs, in vivo vessel forming potential in a naïve or clonal expanded state, clonal proliferative potential, and ECFC self-renewal from recovered implanted vessels in host mice. One might be tempted to conclude identification of a circulating endothelial stem/progenitor cell state, as has been reported for VESP. However, such a claim must prove that putative endothelial stem/progenitor cells can become arteries, veins, capillaries, and lymph vessels and that the cells can repopulate these diverse vascular states after injury (Chambers et al., 2021). Future work will be required to prove murine C-ECFC possess endothelial stem/progenitor potency.

The negligible number of C-ECFCs at E18.5 and their sudden appearance at P0 (Figure 1A) suggests that emergence of TT^+^ colonies in the blood at birth is unlikely to have originated from the pre-existing C-ECFCs at E18.5; such a burst in circulating cells would require an approximate 8-fold increase in < 36h. Given the greater proliferative potential of emerging clones of resident vascular ECFCs (Figure S2C−S2E) it is more plausible that they reside upstream in the hierarchy and would give rise to C-ECFCs. Additionally, we provide clear evidence for naïve C-ECFC possession of in vivo vessel forming potential in mouse and man. These data resolve any lingering challenges that naïve C-ECFC lack vasculogenic activity unless isolated and expanded in culture with added endothelial growth factors. Likewise, lineage tracing results with Flt3cre; mTmG mice have clarified C-ECFC are not derived from bone marrow hematopoietic cells as originally predicted (Asahara *et al*., 1997).

ScRNAseq results have identified several C-ECs subsets in human CB. Of greatest interest, C16 is enriched with TF regulons for EC early development and ECM genes such as fibulin, decorin, and matrix gla-protein that are critical for cell adhesion (Senger and Davis, 2011) and vasculogenesis (Jarvelainen et al., 2015; Maishi and Hida, 2017) (Figure 3F). In the integrated UMAP and trajectory analysis (Figures 3G−3I), C16 localized closely with HUVEC, retained upregulated ECM genes, and existed in the transitional region (HUVEC to CBMNC EC) in pseudotime, suggesting this population was derived from resident vascular ECs; whether C-ECFC are mobilized from all tissue vasculature remains unknown. Since PROCR was the only CD marker enriched in C16, we showed that PROCR^high^ cell fraction and the CD34^bright^ population are clearly enriched in EC colony formation (Figures 4B, 4C and 4E). We conclude the CD34^bright^ and PROCR^high^ C-ECs as displaying properties of C-ECFCs (Figure 4E), providing novel insights that C-ECFCs can be potentially enriched for further studies toward translation to the clinic.

### Limitation of the study

We did not perform transcriptome studies that can explore the potential marker gene expression for murine C-ECFCs given the small amount of blood that can be obtained from each postnatal mouse. Furthermore, we did not prospectively isolate human C-ECFCs for in vivo vessel forming studies and we inferred the CD34^bright^PROCR^+^ population is responsible for EC colony formation based on our sequential magnetic isolation results. Finally, we can only deduce that C-ECFCs are mobilized from resident vessel ECFC through use of several lineage tracing transgenic approaches in mice and scRNAseq in human studies, as we cannot directly image the emergence of C-ECFCs into circulation induced by metabolic or physiologic changes occurring at birth.

## Acknowledgements

We thank Tiffany Lewallen for her administrative assistance, and also thank Emily Sims (Indiana University AngioBioCore) and members of department of obstetrics and gynecology in Nara Medical University for providing the access to umbilical cord and cord blood. The authors are grateful to Slava Epelman (University of Toronto, Toronto, Canada) for the gift of Flt3Cre+;ROSA^mTmG/mTmG^ mice and Yi Zheng (Cincinnati Children’s Hospital Medical Center, Cincinnati, USA) for the gift of Tie2Cre mice. Animal study was approved by the IACUC of the Indiana University School of Medicine. Human study was approved by Nara Medical University Ethics Committee. Funding: AHA Postdoctoral Fellowship 836459 (Y.L.), NYSTEM Training Program in Stem Cell Biology and Regenerative Medicine (Y.L.), Shinya Foundation for International Exchange of Osaka University Graduate School of Medicine Grant (K.B.), JSPS Overseas Research Fellowships (K.B.), AHA 17GRNT33671036 (P.D.B), National Heart, Lung and Blood Institute grant R01HL128827 (M.P.M.), Nara Medical University Grant-in-Aid for Collaborative Research Projects (K.H.), NIH 5R21 CA202296 and 2RO1 EY012601-17 (M.C.Y.).

## Author contributions

Conceptualization; Y.L., K.B., C.H.G., and M.C.Y. Methodology; Y.L., K.B., C.H.G., W.C.S., M.Y., N.P., S.P.T., X.X., R.H.A., K.N., and J.Y. Investigation; K.B., J.M., and T.H. Data curation; H.G. Writing: Y.L., K.B., C.H.G., and M.C.Y. Supervision: Y.L.L, K.B., K.H., and M.C.Y. Funding acquisition: Y.L., K.B., P.D.B, M.P.M., K.H., and M.C.Y.

## Inclusion and diversity

We worked to ensure gender balance in our reference list while citing work relevant to this study.

## References

Aibar, S., Gonzalez-Blas, C.B., Moerman, T., Huynh-Thu, V.A., Imrichova, H., Hulselmans, G., Rambow, F., Marine, J.C., Geurts, P., Aerts, J., et al. (2017). SCENIC: single-cell regulatory network inference and clustering. Nat Methods 14, 1083–1086. 10.1038/nmeth.4463.

Aplin, J.D., Whittaker, H., Jana Lim, Y.T., Swietlik, S., Charnock, J., and Jones, C.J. (2015). Hemangioblastic foci in human first trimester placenta: Distribution and gestational profile. Placenta 36, 1069–1077. 10.1016/j.placenta.2015.08.005.

Asahara, T., Murohara, T., Sullivan, A., Silver, M., van der Zee, R., Li, T., Witzenbichler, B., Schatteman, G., and Isner, J.M. (1997). Isolation of putative progenitor endothelial cells for angiogenesis. Science 275, 964–967.

Bertolini, F., Shaked, Y., Mancuso, P., and Kerbel, R.S. (2006). The multifaceted circulating endothelial cell in cancer: towards marker and target identification. Nat Rev Cancer 6, 835–845. 10.1038/nrc1971.

Blann, A.D., Woywodt, A., Bertolini, F., Bull, T.M., Buyon, J.P., Clancy, R.M., Haubitz, M., Hebbel, R.P., Lip, G.Y., Mancuso, P., et al. (2005). Circulating endothelial cells. Biomarker of vascular disease. Thromb Haemost 93, 228–235. 10.1160/TH04-09-0578.

Boyer, S.W., Schroeder, A.V., Smith-Berdan, S., and Forsberg, E.C. (2011). All hematopoietic cells develop from hematopoietic stem cells through Flk2/Flt3-positive progenitor cells. Cell Stem Cell 9, 64–73. 10.1016/j.stem.2011.04.021.

Case, J., Mead, L.E., Bessler, W.K., Prater, D., White, H.A., Saadatzadeh, M.R., Bhavsar, J.R., Yoder, M.C., Haneline, L.S., and Ingram, D.A. (2007). Human CD34+AC133+VEGFR-2+ cells are not endothelial progenitor cells but distinct, primitive hematopoietic progenitors. Exp Hematol 35, 1109–1118. 10.1016/j.exphem.2007.04.002.

Chambers, S.E.J., Pathak, V., Pedrini, E., Soret, L., Gendron, N., Guerin, C.L., Stitt, A.W., Smadja, D.M., and Medina, R.J. (2021). Current concepts on endothelial stem cells definition, location, and markers. Stem Cells Transl Med 10 Suppl 2, S54–S61. 10.1002/sctm.21-0022.

Estes, M.L., Mund, J.A., Ingram, D.A., and Case, J. (2010). Identification of endothelial cells and progenitor cell subsets in human peripheral blood. Curr Protoc Cytom Chapter 9, Unit 9 33 31–11. 10.1002/0471142956.cy0933s52.

Feaver, R.E., Gelfand, B.D., and Blackman, B.R. (2013). Human haemodynamic frequency harmonics regulate the inflammatory phenotype of vascular endothelial cells. Nat Commun 4, 1525. 10.1038/ncomms2530.

Guervilly, C., Burtey, S., Sabatier, F., Cauchois, R., Lano, G., Abdili, E., Daviet, F., Arnaud, L., Brunet, P., Hraiech, S., et al. (2020). Circulating Endothelial Cells as a Marker of Endothelial Injury in Severe COVID -19. J Infect Dis 222, 1789–1793. 10.1093/infdis/jiaa528.

Hebbel, R.P. (2017). Blood endothelial cells: utility from ambiguity. J Clin Invest 127, 1613–1615. 10.1172/JCI93649.

Hirashima, M., Kataoka, H., Nishikawa, S., Matsuyoshi, N., and Nishikawa, S. (1999). Maturation of embryonic stem cells into endothelial cells in an in vitro model of vasculogenesis. Blood 93, 1253–1263.

Hochstetler, C.L., Feng, Y., Sacma, M., Davis, A.K., Rao, M., Kuan, C.Y., You, L.R., Geiger, H., and Zheng, Y. (2019). KRas(G12D) expression in the bone marrow vascular niche affects hematopoiesis with inflammatory signals. Exp Hematol 79, 3–15 e14. 10.1016/j.exphem.2019.10.003.

Ingram, D.A., Mead, L.E., Moore, D.B., Woodard, W., Fenoglio, A., and Yoder, M.C. (2005). Vessel wall-derived endothelial cells rapidly proliferate because they contain a complete hierarchy of endothelial progenitor cells. Blood 105, 2783–2786. 10.1182/blood-2004-08-3057.

Ingram, D.A., Mead, L.E., Tanaka, H., Meade, V., Fenoglio, A., Mortell, K., Pollok, K., Ferkowicz, M.J., Gilley, D., and Yoder, M.C. (2004). Identification of a novel hierarchy of endothelial progenitor cells using human peripheral and umbilical cord blood. Blood 104, 2752–2760. 10.1182/blood-2004-04-1396.

Jarvelainen, H., Sainio, A., and Wight, T.N. (2015). Pivotal role for decorin in angiogenesis. Matrix Biol 43, 15–26. 10.1016/j.matbio.2015.01.023.

Kawasaki, T., Nishiwaki, T., Sekine, A., Nishimura, R., Suda, R., Urushibara, T., Suzuki, T., Takayanagi, S., Terada, J., Sakao, S., and Tatsumi, K. (2015). Vascular Repair by Tissue-Resident Endothelial Progenitor Cells in Endotoxin-Induced Lung Injury. Am J Respir Cell Mol Biol 53, 500–512. 10.1165/rcmb.2014-0185OC.

Lanuti, P., Rotta, G., Almici, C., Avvisati, G., Budillon, A., Doretto, P., Malara, N., Marini, M., Neva, A., Simeone, P., et al. (2016). Endothelial progenitor cells, defined by the simultaneous surface expression of VEGFR2 and CD133, are not detectable in healthy peripheral and cord blood. Cytometry A 89, 259–270. 10.1002/cyto.a.22730.

Lin, Y., Gil, C.H., and Yoder, M.C. (2019). Identification of Circulating Endothelial Colony-Forming Cells from Murine Embryonic Peripheral Blood. Methods Mol Biol 1940, 97–107. 10.1007/978-1-4939-9086-3_7.

Maishi, N., and Hida, K. (2017). Tumor endothelial cells accelerate tumor metastasis. Cancer Sci 108, 1921–1926. 10.1111/cas.13336.

McDonald, A.I., Shirali, A.S., Aragon, R., Ma, F., Hernandez, G., Vaughn, D.A., Mack, J.J., Lim, T.Y., Sunshine, H., Zhao, P., et al. (2018). Endothelial Regeneration of Large Vessels Is a Biphasic Process Driven by Local Cells with Distinct Proliferative Capacities. Cell Stem Cell 23, 210–225 e216. 10.1016/j.stem.2018.07.011.

Medina, R.J., Barber, C.L., Sabatier, F., Dignat-George, F., Melero-Martin, J.M., Khosrotehrani, K., Ohneda, O., Randi, A.M., Chan, J.K.Y., Yamaguchi, T., et al. (2017). Endothelial Progenitors: A Consensus Statement on Nomenclature. Stem Cells Transl Med 6, 1316–1320. 10.1002/sctm.16-0360.

Mund, J.A., Estes, M.L., Yoder, M.C., Ingram, D.A., Jr., and Case, J. (2012). Flow cytometric identification and functional characterization of immature and mature circulating endothelial cells. Arterioscler Thromb Vasc Biol 32, 1045–1053. 10.1161/ATVBAHA.111.244210.

Novershtern, N., Subramanian, A., Lawton, L.N., Mak, R.H., Haining, W.N., McConkey, M.E., Habib, N., Yosef, N., Chang, C.Y., Shay, T., et al. (2011). Densely interconnected transcriptional circuits control cell states in human hematopoiesis. Cell 144, 296–309. 10.1016/j.cell.2011.01.004.

Purhonen, S., Palm, J., Rossi, D., Kaskenpaa, N., Rajantie, I., Yla-Herttuala, S., Alitalo, K., Weissman, I.L., and Salven, P. (2008). Bone marrow-derived circulating endothelial precursors do not contribute to vascular endothelium and are not needed for tumor growth. Proc Natl Acad Sci U S A 105, 6620–6625. 10.1073/pnas.0710516105.

Ricard, N., Bailly, S., Guignabert, C., and Simons, M. (2021). The quiescent endothelium: signalling pathways regulating organ-specific endothelial normalcy. Nat Rev Cardiol 18, 565–580. 10.1038/s41569-021-00517-4.

Senger, D.R., and Davis, G.E. (2011). Angiogenesis. Cold Spring Harb Perspect Biol 3, a005090. 10.1101/cshperspect.a005090.

Somani, A., Nguyen, J., Milbauer, L.C., Solovey, A., Sajja, S., and Hebbel, R.P. (2007). The establishment of murine blood outgrowth endothelial cells and observations relevant to gene therapy. Transl Res 150, 30–39. 10.1016/j.trsl.2007.02.002.

Tarnawsky, S.P., Yu, W.M., Qu, C.K., Chan, R.J., and Yoder, M.C. (2018). Hematopoietic-restricted Ptpn11E76K reveals indolent MPN progression in mice. Oncotarget 9, 21831–21843. 10.18632/oncotarget.25073.

Tura, O., Skinner, E.M., Barclay, G.R., Samuel, K., Gallagher, R.C., Brittan, M., Hadoke, P.W., Newby, D.E., Turner, M.L., and Mills, N.L. (2013). Late outgrowth endothelial cells resemble mature endothelial cells and are not derived from bone marrow. Stem Cells 31, 338–348. 10.1002/stem.1280.

Wakabayashi, T., Naito, H., Suehiro, J.I., Lin, Y., Kawaji, H., Iba, T., Kouno, T., Ishikawa-Kato, S., Furuno, M., Takara, K., et al. (2018). CD157 Marks Tissue-Resident Endothelial Stem Cells with Homeostatic and Regenerative Properties. Cell Stem Cell 22, 384–397 e386. 10.1016/j.stem.2018.01.010.

Wang, Y., Nakayama, M., Pitulescu, M.E., Schmidt, T.S., Bochenek, M.L., Sakakibara, A., Adams, S., Davy, A., Deutsch, U., Luthi, U., et al. (2010). Ephrin-B2 controls VEGF-induced angiogenesis and lymphangiogenesis. Nature 465, 483–486. 10.1038/nature09002.

Yoder, M.C., Mead, L.E., Prater, D., Krier, T.R., Mroueh, K.N., Li, F., Krasich, R., Temm, C.J., Prchal, J.T., and Ingram, D.A. (2007). Redefining endothelial progenitor cells via clonal analysis and hematopoietic stem/progenitor cell principals. Blood 109, 1801–1809. 10.1182/blood-2006-08-043471.

Zhao, Y., Li, X., Zhao, W., Wang, J., Yu, J., Wan, Z., Gao, K., Yi, G., Wang, X., Fan, B., et al. (2019). Single-cell transcriptomic landscape of nucleated cells in umbilical cord blood. Gigascience 8. 10.1093/gigascience/giz047.

