## Supplemental Figures, Tables, and Methods for "Origin, Prospective Identification, and Function of Circulating Endothelial Colony Forming Cells in Mouse and Human"

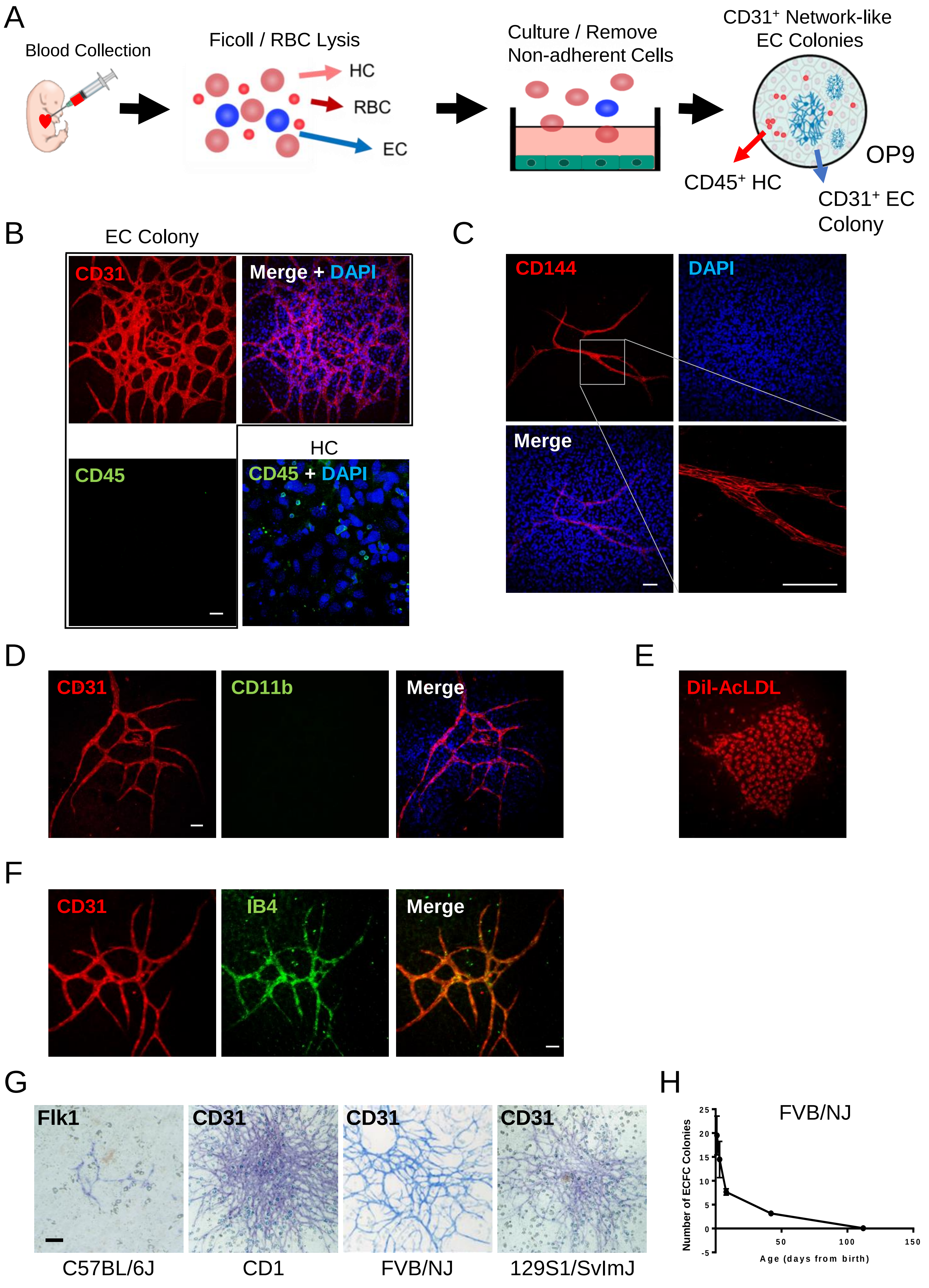
A
Blood Collection
Ficoll / RBC Lysis
HC
RBC
EC
Culture / Remove Non-adherent Cells
CD31+ Network-like EC Colonies
OP9
CD45+ HC
CD31+ EC Colony
B
EC Colony
CD31
Merge + DAPI
CD45
HC
CD45 + DAPI
C
CD144
DAPI
Merge
D
CD31
CD11b
Merge
E
Dil-AcLDL
F
CD31
IB4
Merge
G
Flk1
CD31
CD31
CD31
C57BL/6J
CD1
FVB/NJ
129S1/SvImJ
H
FVB/NJ
Number of ECFC Colonies
Age (days from birth)
25
20
15
10
5
0
-5
50
100
150

**Figure S1. Neonatal/Juvenile Murine Peripheral Blood Contains Circulating Endothelial Colony Forming Cells (C-ECFCs) (Related to Figure 1).**

(A) Schematic of cultured C-ECFC from murine neonatal/juvenile peripheral blood. RBC, red blood cells; HC, hematopoietic cells; EC, endothelial cells. (B) Murine C-ECFC derived EC colonies express the endothelial marker CD31 (red), but not hematopoietic marker CD45 (green). Scale, 100 μm. (C) Murine C-ECFC derived EC colonies express the endothelial marker CD144 (red). Scale, 100 μm. (D) Murine C-ECFC derived EC colonies do not express the myeloid hematopoietic marker CD11b (green). Scale, 100 μm. (E) Murine C-ECFC derived EC colonies can ingest AcLDL (red). Dil-AcLDL, Dil-Acetylated Low Density Lipoprotein. (F) Murine C-ECFC derived EC colonies can bind lectin (green). IB4, isolectin B4. Scale, 100 μm. (G) Mouse C-ECFC derived colonies from the peripheral blood of C57BL/6J, CD1, FVB/NJ and 129S1/SvImJ. Pictures show the staining of alkaline phosphatase conjugated anti-rat IgG secondary antibodies against rat anti-mouse Flk1 (left), or CD31 (other 3 panels). Scale, 200 μm. (H) Kinetics of emergence of C-ECFC derived EC colonies in FVB/NJ mice blood.

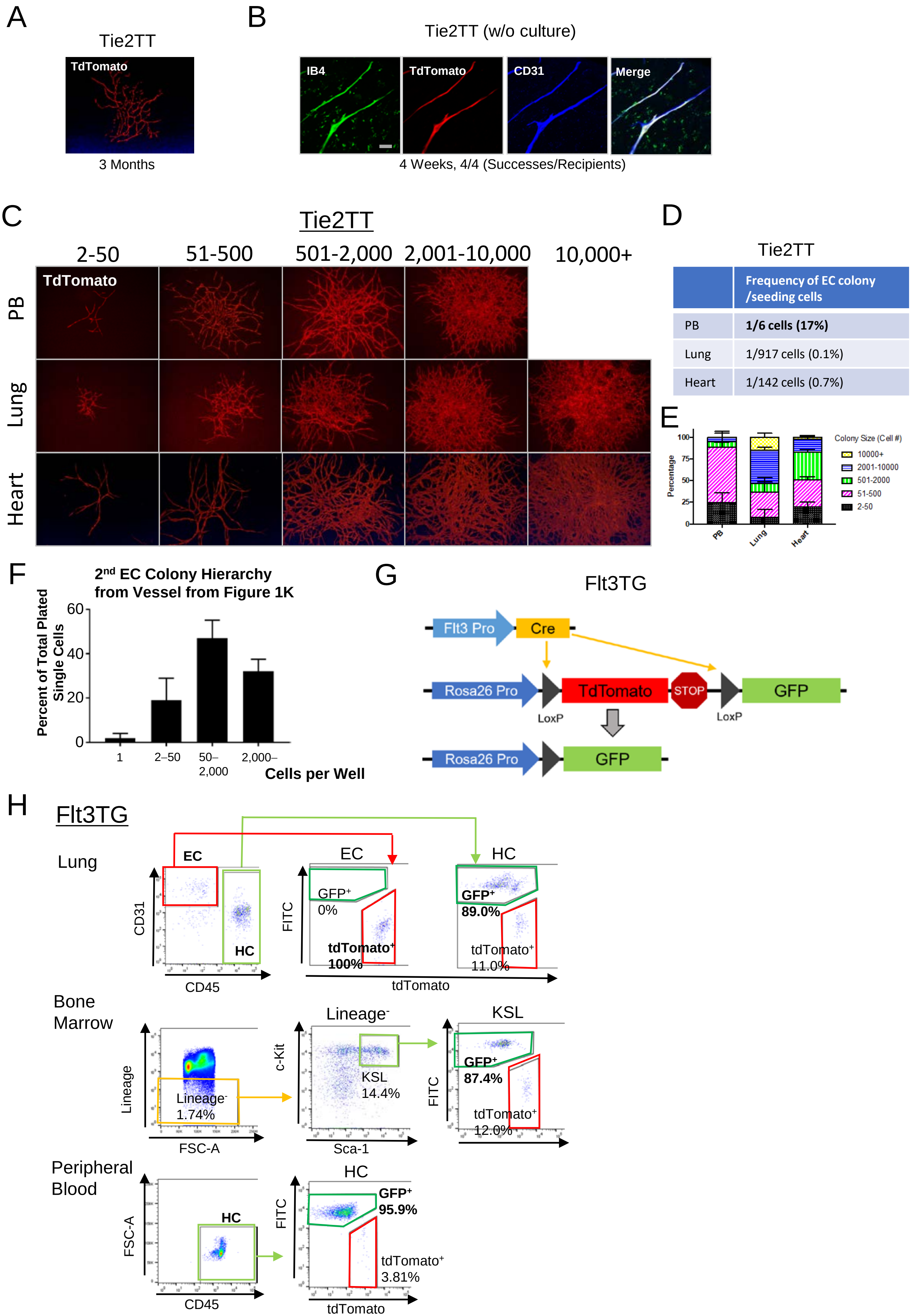

A
Tie2TT
TdTomato
3 Months
B
Tie2TT (w/o culture)
IB4
TdTomato
CD31
Merge
4 Weeks, 4/4 (Successes/Recipients)
C
Tie2TT
2-50
51-500
501-2,000
2,001-10,000
10,000+
PB
Lung
Heart
TdTomato
D
Tie2TT
Frequency of EC colony /seeding cells
PB 1/6 cells (17%)
Lung 1/917 cells (0.1%)
Heart 1/142 cells (0.7%)
E
Percentage
100
75
50
25
0
PB
Lung
Heart
Colony Size (Cell #)
10000+
2001-10000
501-2000
51-500
2-50
F
2nd EC Colony Hierarchy from Vessel from Figure 1K
Percent of Total Plated Single Cells
60
40
20
0
1
2–50
50–2,000
2,000–
Cells per Well
G
Flt3TG
Flt3 Pro
Cre
Rosa26 Pro
TdTomato
STOP
GFP
LoxP
LoxP
Rosa26 Pro
GFP
H
Flt3TG
Lung
EC
HC
CD31
CD45
EC
HC
FITC
GFP+ 0%
tdTomato+ 100%
GFP+ 89.0%
tdTomato+ 11.0%
tdTomato
Bone Marrow
Lineage
FSC-A
Lineage- 1.74%
Lineage-
c-Kit
KSL 14.4%
Sca-1
KSL
FITC
GFP+ 87.4%
tdTomato+ 12.0%
Peripheral Blood
FSC-A
HC
CD45
HC
FITC
GFP+ 95.9%
tdTomato+ 3.81%
tdTomato

**Figure S2. Murine and Human CECs Contain Endothelial Colony Forming Cell That Can Form Functional Blood Vessel in vitro and in vivo, and Schematic and Flow Cytometry Result of Flt3mTmG Mouse for Investigating Their Origin (Related to Figure 1 and Figure 2).**

(A) A representative picture of a $TdTomato^+$ secondary colony from $TdTomato^+$ vessels in the gel of 3 months after transplantation. (B) Four weeks after collagen-plug transplantation using PB of Tie2TT (P2), uncultured CEC derived blood vessels ($TdTomato^+$) are inosculated with host vasculatures (shown by the labeling of isolectin B4). Scale, 50 μm. (C) Representative EC colony images according to the number of cells per colony from PB, lung, heart in Tie2TT mouse (P6). From left to right: 2-50, 51-500, 501-2,000, 2,001-10,000, >10,000 cells per colony are shown. Note that colonies of >10,000 cells can only be detected from lung and heart. (D) Table showing frequency of appearance of EC colonies per seeded cells ($CD34^+CD45^-$ $TdTomato^+$ Fraction); EC colonies from PB are more frequent than from lung or heart. (E) Bar graph showing the percentage of cells per EC colony; PB-derived EC colonies have fewer cells per colony. (F) $2^{nd}$ EC culture grew from human cord blood CEC derived blood vessels show a hierarchy of proliferative potential in single cell colony forming assay. (G) Schematic of Cre-loxP recombination in Flt3Cre;mTmG mouse. Cre recombinase is expressed under the control of Flt3 regulatory elements. Cre-mediated excision of TdTomato results in induction of GFP expression. (H) Flow cytometry analysis of postnatal day 14 Flt3Cre:mTmG mice. GFP did not label EC in the lung but labeled almost all hematopoietic cells and hematopoietic stem cells in the lung, bone marrow and peripheral blood.

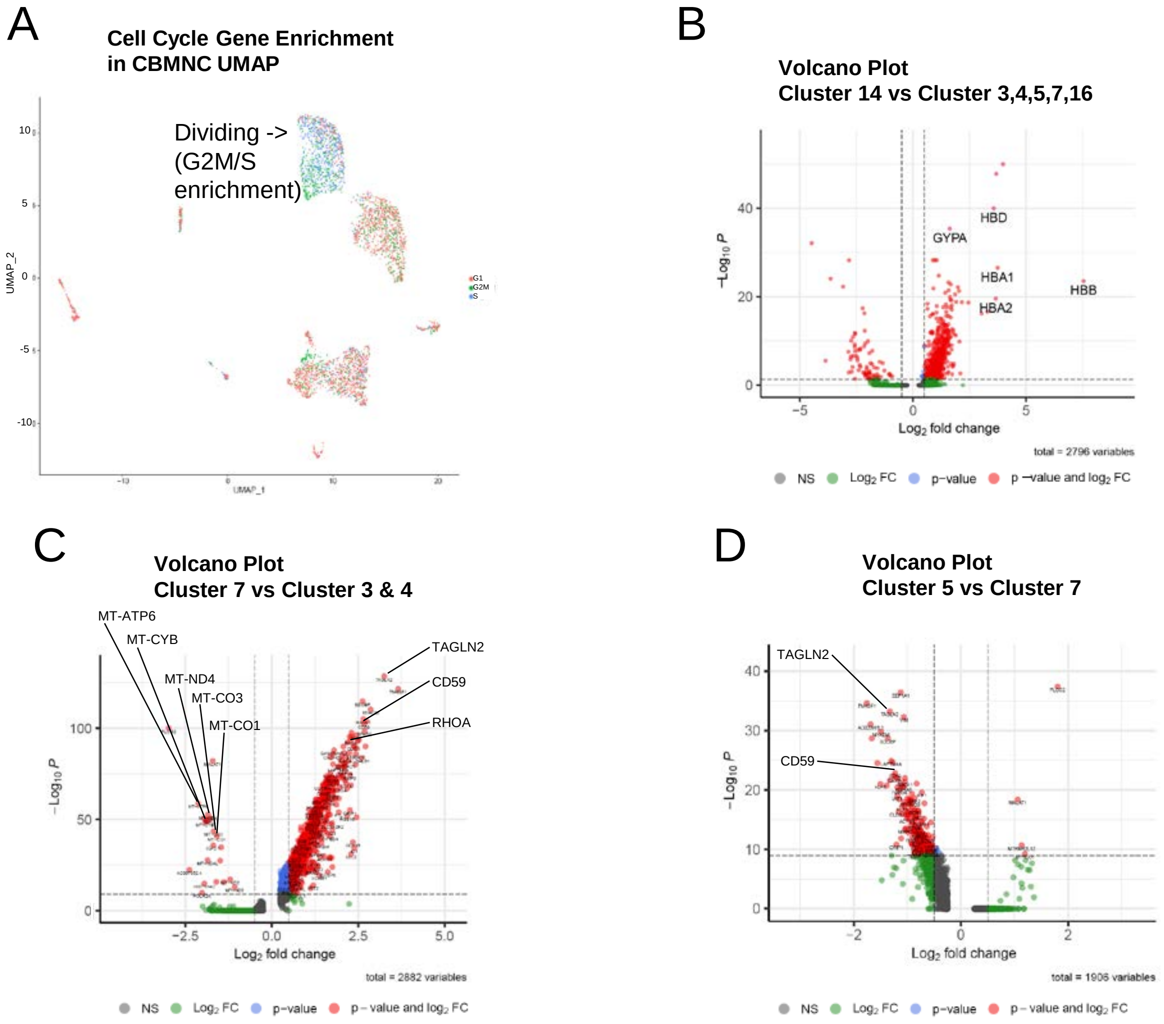

A
Cell Cycle Gene Enrichment in CBMNC UMAP
Dividing -> (G2M/S enrichment)
UMAP_2
UMAP_1
10
5
0
-5
-10
-10
0
10
20
G1
G2M
S
B
Volcano Plot
Cluster 14 vs Cluster 3,4,5,7,16
HBD
GYPA
HBA1
HBB
HBA2
−Log10 P
Log2 fold change
40
20
0
−5
0
5
total = 2796 variables
NS
Log2 FC
p−value
p −value and log2 FC
C
Volcano Plot
Cluster 7 vs Cluster 3 & 4
MT-ATP6
MT-CYB
MT-ND4
MT-CO3
MT-CO1
TAGLN2
CD59
RHOA
−Log10 P
Log2 fold change
100
50
0
−2.5
0.0
2.5
5.0
total = 2882 variables
NS
Log2 FC
p−value
p − value and log2 FC
D
Volcano Plot
Cluster 5 vs Cluster 7
TAGLN2
CD59
−Log10 P
Log2 fold change
40
30
20
10
0
−2
0
2
total = 1906 variables
NS
Log2 FC
p−value
p − value and log2 FC

**Figure S3. Single Cell RNA Sequencing Reveals Unique Endothelial Cell Related Clusters in Circulating Human Cord Blood Mononuclear Cells (Related to Figure 3).**

(A) Cell cycle gene enrichment analysis in UMAP plots of CBMNCs. G2M and S phase specific genes are enriched in cluster 1, 2, and 6 of CBMNCs. (B) Volcano plot comparing cluster 14 vs cluster 3, 4, 5, 7, and 16. Dots are colored based on log2 fold change and p-value. (C) Volcano plot comparing cluster 7 vs cluster 3 &4. (D) Volcano plot comparing cluster 5 and cluster 7.

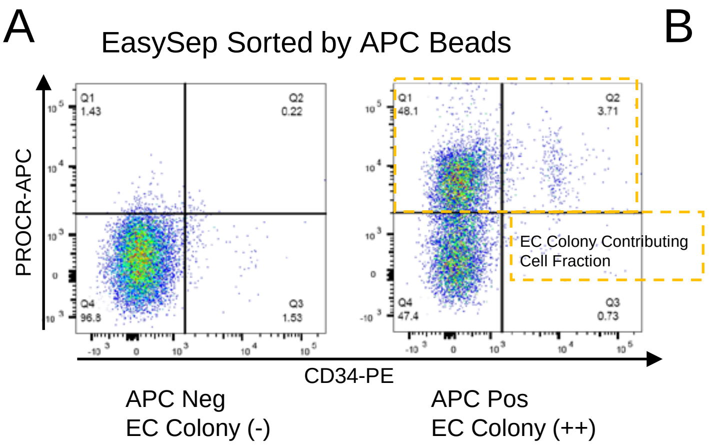


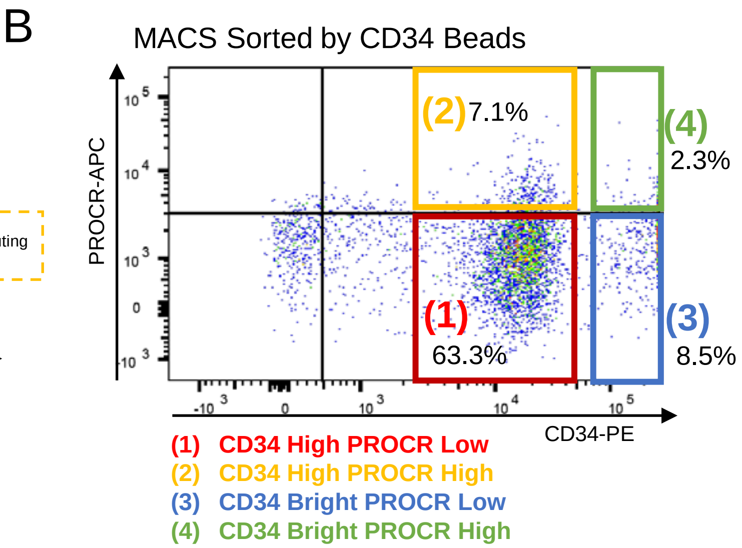


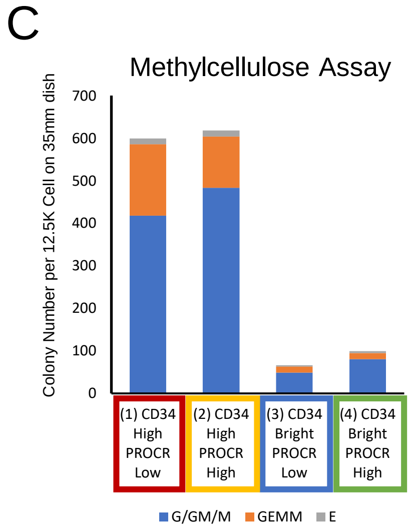


D

In vitro EC Colony-Forming Assay after FACS Sorting

| | Exp.1 | Exp.2 | Exp.3 |
|---|---|---|---|
| (1) CD34 High PROCR Low (216K,175K,268K in each Exp) | 0 | 0 | 0 |
| (2) CD34 High PROCR High (38K,27K,80K in each Exp) | 0 | 0 | 0 |
| (3) CD34 Bright PROCR Low (10K,27K,23K in each Exp) | 0 | 0 | 0 |
| (4) CD34 Bright PROCR High (10K,27k,23K in each Exp) | **3 (ECFC)** | 0 | 0 |

E

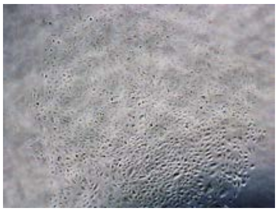

Representative EC Colony (from Exp.1 of Figure S4D)

**Figure S4 (Related to Figure 4 and Table S2).**

(A) Representative flow cytometry analysis of EasySep-sorted CBMNCs by PROCR-APC and anti-APC beads. Cells from APC negative fraction do not yield EC colonies, whereas those from APC positive fraction yield EC colonies. (B) Representative flow cytometry analysis of MACS-sorted CBMNCs by CD34 beads. CD34 positive cells can be divided into 4 fractions using CD34 and PROCR expression levels; (1) CD34 High PROCR High (in red), (2) CD34 High PROCR Low (in orange), (3) CD34 Bright (= very high) PROCR Low (in blue), and (4) CD34 Bright PROCR High (in green). CD34 bright cells represent about 1/8 of total CD34 positive cells. (C) Hematopoietic colony number after methylcellulose assay using cells from 4 fractions sorted by FACS. CD34 high fractions ((1) and (2)) give rise to 6-10 times more hematopoietic colonies compared with CD34 bright fractions ((3) and (4)). (D) Colony numbers after in vitro EC colony-forming assay from 4 fractions sorted by FACS. Only CD34 bright PROCR high fractions (4) yielded EC colonies once. Other fractions did not yield EC colonies. (E) Representative EC colony (about 1,000 cells per colony) with cobblestone appearance from CD34 bright PROCR high fraction (4).

**Table S1. Enriched Regulons by SCENIC Analysis**

**Cluster 16 vs cluster 3, 4, 5, and 7.**

| Regulon (gene number) | Enriched Cluster(s) |
|---|---|
| GABPA (1352) | 16 |
| CREB3 (15) | 16 |
| MAZ_extended (2768) | 16 |
| NR2F2_extended (10) | 16 |
| FOXC2_extended (77) | 16 |
| ZNF358_extended (649) | 16 |
| ATF4 (16) | 16 |
| YY1 (1233) | 16 |
| NFIC_extended (14) | 16 |
| HOXB2_extended (23) | 16 |
| CUX1 (12) | 16 |
| MAX_extended (2634) | 16 |
| FOXC1_extended (24) | 16 |
| NR4A1_extended (16) | 16 |
| ZNF664 (39) | 16 |

| | |
|---|---|
| NFKB2 (72) | 3_4_5_7 |
| XBP1 (237) | 3_4_5_7 |
| JUND (38) | 3_4_5_7 |
| CREM (43) | 3_4_5_7 |
| TAF7 (14) | 3_4_5_7 |
| BCL3_extended (78) | 3_4_5_7 |
| IRF1 (579) | 3_4_5_7 |
| ATF3 (2030) | 3_4_5_7 |
| KLF3_extended (126) | 3_4_5_7 |
| KLF2_extended (92) | 3_4_5_7 |
| JUNB (21) | 3_4_5_7 |
| KLF10 (41) | 3_4_5_7 |
| ATF5 (50) | 3_4_5_7 |
| JUN (29) | 3_4_5_7 |
| KLF4 (236) | 3_4_5_7 |

**Table S1. Enriched Regulons by SCENIC Analysis**

**Cluster 14 vs cluster 3, 4, 5, and 7.**

| Regulon (gene number) | Enriched Cluster(s) |
|---|---|
| REST_extended (657) | 14 |
| YY1 (2148) | 14 |
| SRF (28) | 14 |
| HDAC2_extended (1631) | 14 |
| MAFF_extended (12) | 14 |
| MXD3_extended (49) | 14 |
| BHLHE40_extended (1970) | 14 |
| SAP30_extended (452) | 14 |
| NFIL3 (29) | 14 |
| TCF3 (35) | 14 |
| HOXA10 (13) | 14 |
| MAZ_extended (2751) | 14 |
| NFYB (17) | 14 |
| THAP11 (160) | 14 |
| E2F4 (24) | 14 |
| RAD21_extended (2471) | 14 |
| TFDP1_extended (70) | 14 |
| ZNF664 (38) | 14 |
| HMGN3_extended (11) | 14 |
| TFDP2_extended (35) | 14 |
| TBP_extended (33) | 14 |
| MAX_extended (2734) | 14 |
| POLE3_extended (757) | 14 |

| | |
|---|---|
| NFE2L2 (27) | 3_4_5_7 |
| IRF1 (659) | 3_4_5_7 |
| CREB3_extended (541) | 3_4_5_7 |
| ATF3 (2460) | 3_4_5_7 |
| JUND (41) | 3_4_5_7 |
| TAF7 (28) | 3_4_5_7 |
| XBP1 (192) | 3_4_5_7 |
| ATF4 (40) | 3_4_5_7 |

| | |
|---|---|
| JUNB (24) | 3_4_5_7 |
| ATF5 (136) | 3_4_5_7 |
| KLF10 (94) | 3_4_5_7 |
| KLF2_extended (43) | 3_4_5_7 |
| KLF4 (258) | 3_4_5_7 |
| JUN (29) | 3_4_5_7 |
| NFKB2 (102) | 3_4_5_7 |
| BCL3 (18) | 3_4_5_7 |
| ETS2 (2037) | 3_4_5_7 |
| CREM (108) | 3_4_5_7 |
| CREB5 (58) | 3_4_5_7 |
| CEBPB (50) | 3_4_5_7 |
| DDIT3 (55) | 3_4_5_7 |
| FOSL2 (25) | 3_4_5_7 |
| EGR1_extended (22) | 3_4_5_7 |
| MAFB_extended (28) | 3_4_5_7 |
| ZNF143_extended (21) | 3_4_5_7 |
| GTF2B (11) | 3_4_5_7 |
| SOX4_extended (17) | 3_4_5_7 |
| IRF7 (175) | 3_4_5_7 |
| FOSB_extended (45) | 3_4_5_7 |
| FOS_extended (595) | 3_4_5_7 |
| POLR2A (1612) | 3_4_5_7 |
| IRF9 (94) | 3_4_5_7 |
| IRF2 (155) | 3_4_5_7 |
| EZH2 (33) | 3_4_5_7 |
| FOXK2_extended (48) | 3_4_5_7 |
| POU3F1_extended (21) | 3_4_5_7 |
| PLAGL1 (12) | 3_4_5_7 |
| HOXA13_extended (25) | 3_4_5_7 |
| TAL1_extended (195) | 3_4_5_7 |
| FOXO1_extended (199) | 3_4_5_7 |
| STAT1 (240) | 3_4_5_7 |
| THAP1 (575) | 3_4_5_7 |
| HIST1H2BN_extended (683) | 3_4_5_7 |

| | |
|---|---|
| KDM5B_extended (854) | 3_4_5_7 |
| MYC_extended (2754) | 3_4_5_7 |
| SP3_extended (1034) | 3_4_5_7 |
| RCOR1_extended (724) | 3_4_5_7 |
| MAFK_extended (1157) | 3_4_5_7 |
| ELK4_extended (1112) | 3_4_5_7 |
| ESRRA_extended (2139) | 3_4_5_7 |
| ZNF274_extended (1448) | 3_4_5_7 |
| JDP2 (34) | 3_4_5_7 |
| ATF6_extended (182) | 3_4_5_7 |
| ZBTB7A_extended (162) | 3_4_5_7 |
| STAT2 (314) | 3_4_5_7 |
| FOXO3_extended (81) | 3_4_5_7 |
| SPI1 (257) | 3_4_5_7 |
| EP300 (98) | 3_4_5_7 |
| FOXP1 (87) | 3_4_5_7 |
| FOXC2_extended (178) | 3_4_5_7 |
| ZNF358 (435) | 3_4_5_7 |
| ZNF768 (288) | 3_4_5_7 |
| ZNF91_extended (161) | 3_4_5_7 |
| GABPB1 (443) | 3_4_5_7 |
| USF1_extended (494) | 3_4_5_7 |
| E2F6_extended (471) | 3_4_5_7 |
| ATF6B_extended (255) | 3_4_5_7 |
| GATA2 (15) | 3_4_5_7 |
| MXI1 (372) | 3_4_5_7 |
| ELF2 (830) | 3_4_5_7 |
| BCLAF1_extended (2559) | 3_4_5_7 |
| KDM5A_extended (1564) | 3_4_5_7 |
| IRF3_extended (1223) | 3_4_5_7 |
| SREBF2_extended (844) | 3_4_5_7 |
| TAF1 (639) | 3_4_5_7 |
| SREBF1 (64) | 3_4_5_7 |
| CREB1 (12) | 3_4_5_7 |
| SMARCA4_extended (1182) | 3_4_5_7 |

| | |
|---|---|
| HCFC1_extended (1143) | 3_4_5_7 |
| GABPA (1291) | 3_4_5_7 |
| CTCF_extended (996) | 3_4_5_7 |
| ELF4_extended (618) | 3_4_5_7 |
| ETV6 (31) | 3_4_5_7 |
| PML (1286) | 3_4_5_7 |
| GTF2F1_extended (1461) | 3_4_5_7 |
| FLI1 (12) | 3_4_5_7 |
| ERG_extended (1390) | 3_4_5_7 |
| HDAC6_extended (432) | 3_4_5_7 |
| ETV3 (894) | 3_4_5_7 |
| RELA (540) | 3_4_5_7 |
| ZMIZ1_extended (183) | 3_4_5_7 |
| NR3C1 (560) | 3_4_5_7 |
| ETS1 (1844) | 3_4_5_7 |
| ELK3 (2129) | 3_4_5_7 |
| CEBPZ_extended (1518) | 3_4_5_7 |
| ELF1 (1897) | 3_4_5_7 |
| ZEB1_extended (136) | 3_4_5_7 |
| SP1 (451) | 3_4_5_7 |
| PATZ1_extended (19) | 3_4_5_7 |
| FOXC1 (11) | 3_4_5_7 |
| TGIF2 (12) | 3_4_5_7 |
| NFIA (10) | 3_4_5_7 |
| ARID3A (23) | 3_4_5_7 |
| MEF2C_extended (11) | 3_4_5_7 |
| SNAI1 (10) | 3_4_5_7 |
| MAFG_extended (11) | 3_4_5_7 |
| KLF11_extended (15) | 3_4_5_7 |
| RORA_extended (11) | 3_4_5_7 |
| NR2F6 (12) | 3_4_5_7 |
| TGIF1 (19) | 3_4_5_7 |
| NFKB1 (13) | 3_4_5_7 |
| FOXN2 (24) | 3_4_5_7 |
| CEBPG (11) | 3_4_5_7 |

| | |
|---|---|
| PRDM1_extended (13) | 3_4_5_7 |
| MLX_extended (12) | 3_4_5_7 |
| ATF7_extended (22) | 3_4_5_7 |
| KLF3_extended (58) | 3_4_5_7 |
| KLF13_extended (31) | 3_4_5_7 |
| NR1H2_extended (25) | 3_4_5_7 |
| NR4A1_extended (12) | 3_4_5_7 |
| ZNF335 (30) | 3_4_5_7 |
| MEF2A_extended (17) | 3_4_5_7 |
| NR2F2_extended (31) | 3_4_5_7 |
| KLF9_extended (25) | 3_4_5_7 |
| TFE3_extended (34) | 3_4_5_7 |
| TBL1XR1 (12) | 3_4_5_7 |
| MEF2D_extended (12) | 3_4_5_7 |
| ARNT (43) | 3_4_5_7 |
| RFX3_extended (17) | 3_4_5_7 |
| ERF (16) | 3_4_5_7 |
| RXRA_extended (31) | 3_4_5_7 |
| MXD4_extended (68) | 3_4_5_7 |
| NFATC2_extended (25) | 3_4_5_7 |
| ZNF121 (70) | 3_4_5_7 |
| TCF12_extended (20) | 3_4_5_7 |
| TEAD2_extended (11) | 3_4_5_7 |
| ATF2_extended (25) | 3_4_5_7 |
| TBX2_extended (19) | 3_4_5_7 |
| FOXJ3_extended (18) | 3_4_5_7 |
| UBTF (17) | 3_4_5_7 |
| UBP1_extended (40) | 3_4_5_7 |
| NR1H3_extended (10) | 3_4_5_7 |
| ZNF134 (29) | 3_4_5_7 |
| PBX1 (15) | 3_4_5_7 |
| STAT3_extended (11) | 3_4_5_7 |
| RARA_extended (26) | 3_4_5_7 |
| KAT2A_extended (11) | 3_4_5_7 |
| SP6_extended (19) | 3_4_5_7 |

| | |
|---|---|
| SMAD3_extended (11) | 3_4_5_7 |
| ZNF493 (16) | 3_4_5_7 |
| NFYC_extended (38) | 3_4_5_7 |
| NR2C2_extended (13) | 3_4_5_7 |
| HIVEP2_extended (29) | 3_4_5_7 |
| ZNF217_extended (11) | 3_4_5_7 |
| BACH1 (13) | 3_4_5_7 |
| AHR (14) | 3_4_5_7 |

**Table S1. Enriched Regulons by SCENIC Analysis (Related to Figure 3).**

Significantly enriched regulons in indicated clusters were shown.

**Table S2. In vitro EC Colony-Forming Assay Results after Magnetic Sorting**

**MACS or EasySep Sort PROCR-APC & Anti-APC Beads (Figure 4B and Figure S4A)**

| | Exp.1 | Exp.2 | Exp.3 |
|---|---|---|---|
| PROCR-APC High (MACS) (100K in every Exp) | 9 | 18 | 30 |
| PROCR-APC Low (MACS) (1M, 1M, 500K in each Exp) | 0 | 0 | 0 |
| PROCR-APC High (EasySep) (250K in every Exp) | 2 | 13 | Not Performed |
| PROCR-APC Low (EasySep) (1M, 500K in each Exp) | 0 | 0 | Not Performed |

**MACS CD34 MultiSort Beads > Beads Releasing Reagent > MACS PROCR-APC & Anti-APC Beads (Figure 4C)**

| | Exp.1 | Exp.2 | Exp.3 | Exp.4 |
|---|---|---|---|---|
| CD34 Positive > Released > PROCR-APC High (200K, 75K, 250K, 140K in each Exp) | 0 | 0 | 0 | 0 |
| CD34 Positive > Released > PROCR-APC Low (400K, 300K, 500K, 280K in each Exp) | 0 | 0 | 0 | 0 |
| CD34 Positive > Unreleased (75K, 125K, 70K in each Exp) | Not Performed | 4 | 20 | 18 |

**Table S2. In vitro EC Colony Forming Assay Results after FACS Sorting (Related Figure 4 and Figure S4).**

EC colony number is shown in each experiment (Exp). Magnetic sorting with MACS or EasySep is performed before seeding cells on collagen-coated 24 well plate. Detailed magnetic sorting procedures are indicated in Figure 4A. Indicated cell number with parentheses in order was seeded and colonies were evaluated after 2-week culture with EGM-2. Only HPP (high proliferative-potential)-, LPP (low proliferative-potential)-, and cluster ECFCs were counted on day 7-14.

**Table S3. Primers**

**qPCR Primers (Human)**

| | |
|---|---|
| ACTB fwd | CCAACCGCGAGAAGATGA |
| ACTB rev | CCAGAGGCGTACAGGGATAG |
| CD34 fwd | TCCAGAGACAACCTTGAAGC |
| CD34 rev | CTTCTTAAACTCCGCACAGC |
| CDH5 fwd | AGACCACGCCTCTGTCATGTACCAAATC |
| CDH5 rev | CACGATCTCATACCTGGCCTGCTTC |
| FLI1 fwd | AGCGTTAGCAAATGCAGCAAGCTGGT |
| FLI rev | ATTGCCTCACATGCTCCTGTGTCCA |
| GATA2 fwd2 | AGACGACAACCACCACCTTA |
| GATA2 rev2 | TCCTTCTTCATGGTCAGTGG |
| MCAM fwd | AACACAGTGGGCGCTATGAA |
| MCAM rev | AACTCGAGGTCCTGGCTACT |
| PECAM1 fwd | GGTCAGCAGCATCGTGGTCAACATAAC |
| PECAM1 rev | TGGAGCAGGACAGGTTCAGTCTTTCA |
| PROCR fwd | CCAACACCACGATCATTCAG |
| PROCR rev | ATACCGAGTGCGGTTGTAGG |
| PROM1 fwd | CCTCTGGTGGGGTATTTCTTT |
| PROM1 rev | CCAGTTTCCGACTCCTTTTG |
| PTPRC fwd | TAGGGACACGGCTGACTTCCAGATATGA |
| PTPRC rev | GTGTTGGGCTTTGCCCTGTCACAAATAC |

**Genotyping Primers (Mouse)**

| | |
|---|---|
| Cre fwd | CGGTCGATGCAACGAGTGAT |
| Cre rev | CCACCGTCAGTACGTGAGAT |
| Rosa fwd | CTGTTCCTGTACGGCATGG |
| Rosa rev | GGCATTAAAGCAGCGTATCC |

**STAR Methods**

**Animals**

All animal experiments were conducted in accordance with the Guidelines for the Care and Use of Laboratory Animals, and all protocols were approved by Institutional Animal Care and Use Committee of the Indiana University School of Medicine. C57BL/6J (JAX stock #000664), FVB/NJ (FVB, JAX stock #001800), 129S1/SvImJ (SV129, JAX stock #002448), B6.Cg-*Gt(ROSA)26Sor*$^{tm14(CAG\text{-}tdTomato)Hze}$/J (Madisen et al., 2010) (R26R-TdTomato, JAX stock #007914), NOD.Cg-*Prkdc*$^{scid}$/J (NOD/SCID, JAX stock #001303) were purchased from the Jackson Laboratory. CD1 mice (#022) were purchased from Charles River Laboratories. Flt3Cre+;ROSA$^{mTmG/mTmG}$ mice (Benz et al., 2008; Epelman et al., 2014; Muzumdar et al., 2007) were a kind gift from Dr. Slava Epelman, University of Toronto. Tie2CreERT2 mice (kind gift of Dr. Ye Zheng, University of Cincinnati) previously described (Hochstetler et al., 2019) were crossed with the commercially available R26R-TdTomato mice (above) to generate (Tie2TT) mice. The primers used for genotyping the above mentioned Cre or ROSA mice are shown in Table S3. To induce Cre expression in Tie2CreERT2 and Cdh5(PAC)-CreERT2 mice, 50 mg/kg tamoxifen was injected into the animals intra-peritoneally (i.p.) at appropriate time points (3-5 days).

**Murine cell collection**

Blood was collected from postnatal mice by cardiac puncture. After blood collection, mononuclear cells were isolated by re-suspending in red blood cell lysis buffer (Qiagen) for 10 minutes (for culture) or density gradient centrifugation using Histopaque-1083 (Sigma) (for flow cytometry).

To collect cells from mouse lung, liver or heart, tissues were dissected from euthanized mice and were minced with razor blades. Samples were digested with 0.25% collagenase I (Stem Cell Technologies) at 37°C for 30 minutes. After digestion, the samples were re-suspended in medium, pipetted thoroughly, and passed through 70 μm cell strainers to removed cell clumps. To collect mouse bone marrow cells, tibias and femurs were dissected and cleaned with scissors to remove remaining muscle tissues. Then the bones were crushed with a pestle and mortar before the cells were digested and filtered like the other tissues above.

**Human cell collection**

Human umbilical cords and umbilical cord blood samples, destined to be discarded (nonidentified surgical waste and not considered human research) were collected immediately after Cesarean section in Indiana University Health Methodist Hospital.

Human umbilical cord blood samples were also collected in Nara Medical University after informed consent, approved by Nara Medical University Ethics Committee (No. G151).

To collect human umbilical cord vein ECs (HUVECs), the umbilical vessels were flushed with PBS 3 times. Then one end of the vessel was clamped and liberase solution (Roche, 500 μl stock solution diluted with 24.5 ml PBS) was infused into the vessel through the open end before it was clamped. The liberase infused vascular tissues were incubated at 37°C for 14 minutes to release ECs from the basement membrane. Finally, the solution containing digested ECs was flushed into 50 ml tubes for centrifugation and EC recovery.

To collect mononuclear cells from human umbilical cord blood samples, Ficoll-Paque (GE Healthcare) was added to the anti-coagulated blood (diluted in PBS) and cell separation was performed according to the manufacturer's protocol.

**Magnetic activated cell sorting (MACS) for single cell RNAseq (scRNAseq)**

Cord blood mononuclear cells (CBMNCs) and HUVECs for scRNAseq were MACS sorted with CD45-, CD235a-, and CD34-MicroBeads (Miltenyi) using the following method. First, freshly isolated CBMNCs and HUVECs were resuspended in PBS buffer (PBS with 10% FBS (Hyclone)). Then, 10-20 μl of CD45 MicroBeads (human, 130-045-801) and 10-

20 μl CD235a (Glycophorin A) MicroBeads (human, 130-050-501) were added per $10^7$ total cells and incubated at 4°C for 13 minutes (and tubes were tapped every 3-4 minutes). Cells were washed, centrifuged at 400 g for 7 minutes, and resuspended $10^8$ cells in 500 μl of PBS buffer. Cell suspensions were then applied to LD column. Unlabeled cells ($CD45^-CD235a^-$ enriched fraction) which passed through the column were collected and counted for the next step. Collected cells were centrifuged at 400 g for 5 minutes, resuspended with 60 μl of PBS buffer, and 20 μl of Fc Block reagent and 20 μl CD34 Microbeads (human, 130-046-702) were added and incubated at 4°C for 20 minutes (and tubes were tapped every 3-4 minutes). Cells were washed and centrifuged at 400 g for 5 minutes, resuspended in 500 μl of PBS buffer. Cell suspensions were then applied to LS column. After washing 3 times, magnetically labeled cells suspensions were immediately flushed out, filtered with 30 μm filter (Sysmex), and counted. $CD45^-CD235a^-CD34^+$ enriched CBMNCs and HUVECs ("scRNAseq samples") were then suspended in DMEM (Gibco)/10%FBS (Hyclone) in 500-1,000 cells per 1 μl concentration, and used for scRNAseq (see below).

**Single cell library preparation**

The "scRNAseq samples" (freshly isolated and MACS sorted CBMNC and freshly isolated and MACS sorted HUVEC from a same individual human subject) were applied to a single cell master mix with lysis buffer and reverse transcription reagents, following the Chromium Single Cell 3' Reagent Kits V2 and V3 User Guide (10x Genomics). This was followed by cDNA synthesis and library preparation. All libraries were sequenced in Illumina NovaSeq6000 platform in paired-end mode (28 bp + 91 bp). The total number of CBMNC and HUVEC were 4,477 cells and 13,651 cells, respectively. 85K reads per cells were generated and 94% of the sequencing reads reached Q-score at least 30 (Q30) in CBMNC, while 28K reads per cells were generated and 95% of the sequencing reads reached Q30 in HUVEC.

**Single cell data processing**

The 10x Genomics Cell Ranger (v. 2.1.0) pipeline was utilized to demultiplex raw base call files to FASTQ files and reads aligned to the human reference genome GRCh38 using RNAseq aligner STAR (Dobin et al., 2013). Cell Ranger computational output was then analyzed in R (v.3.5.0) using the Seurat package v. 3.0.1 (Stuart et al., 2019). Seurat objects were created for non-integrated and integrated data using the following filtering metrics: gene counts were set between 200–3,000 and mitochondrial gene percentages less than 25 in CBMNCs and HUVEC, gene counts were set between 3,500-10,000 and mitochondrial gene percentages less than 10 to exclude doublets and poor-quality cells. We then removed ribosomal protein genes to reduce noise of dead/dying cells, Gene counts were log transformed and scaled to 0-5. The top 20 principal components were used to perform unsupervised clustering analysis, and visualized using UMAP dimensionality reduction (resolution 0.8). Using the Seurat package, annotation and grouping of clusters to cell type was performed manually by inspection of differentially expressed genes (DEGs) for each cluster, based on canonical marker genes in the literature (Novershtern et al., 2011; Zhao et al., 2019).

**Upstream regulatory network analysis and trajectory analysis**

SCENIC analysis (Aibar et al., 2017) was performed using the default setting and hg19-500bp-upstream-7species.mc9nr.feather database was used for data display.

**Pseudotemporal ordering of single cells**

We performed pseudotime analysis on the integrated Seurat object containing all cells in HUVEC clusters without C12 as well as those in EC clusters (C3, 4, 5, 7, 14, and 16). The datasets were analyzed through the R package Monocle using default parameters. Outputs were obtained detailing the pseudotime cell distributions for each cell type (including cells in cluster 16, which were colored in red). Positional information for the monocle plot was used to color cells (Trapnell et al., 2014).

**Magnetic enrichment of CD34 bright, CD34 high fractions, PROCR high and low fractions.**

To enrich PROCR high and low fractions (Figures 4B), freshly isolated CBMNCs were magnetically sorted using CD201 (EPCR) Antibody (Miltenyi, clone REA337) and Anti-APC MicroBeads (Miltenyi, 130-090-855) or EasySep APC Positive Selection Kit (Stemcell Technologies, ST-18453) according to each manufacturer's protocol.

To enrich the CD34$^{high}$PROCR$^{high}$ fraction, we initially planned to perform a two-step MACS using CD34 MultiSort MicroBeads (Miltenyii). The kit can separate CD34 antibody (Ab) bound to the cell surface from the conjugated magnetic beads by using a proprietary enzyme called "release reagent" after sorting with the Ab-beads tagged cells in the first step. This allows for a second-step Ab-beads tagged reaction for isolating PROCR antibody bound cells. However, we noticed that there were cells in a fraction (unreleased fraction) whose beads could not be separated by the release reagent. This fraction may be caused by the fact that the amount of CD34 antigen on the cell surface is very high (i.e., CD34$^{bright}$), therefore, not all of the Ab-bead tagged cells can be separated in the prescribed release reagent reaction (Figure 4A). Therefore, we proceeded to isolate cells in the following manner.

Freshly isolated cord blood mononuclear cells were counted and labeled with 100 μl of Fc block reagent and 100 μl of CD34 MultiSort MicroBeads (human, 130-056-701) per $10^8$ total cells. Then, the first magnetic separation was performed with LS columns. After magnetically labeled CD34 positive fraction was flushed out, 20 μl of MultiSort Release Reagent was added per 1 mL of cell suspension and incubated for 10 minutes in the refrigerator in the dark (and tubes were tapped every 3-4 minutes). Next, the second magnetic separation was performed. This column can bind bead-unreleased cells; $CD34^{bright}$ cells. Consequently, the magnetic (unreleased) cell fraction was enriched with $CD34^{bright}$ cells and non-magnetic (released) cell fraction was enriched with $CD34^{high}$ cells. For non-magnetic (released) cell fraction, a third separation was able to be performed to sort PROCR high and low fractions as described above (Figures 4A and 4C). The efficiency of separation was evaluated by flow cytometry. All cell fractions were then assessed for proliferative potential by endothelial colony forming culture.

**Flow cytometry**

The following anti-mouse antibodies conjugated with different fluorochrome were used for flow cytometry sorting and analysis: CD31 (clone 390), CD45 (30-F11), Ter119 (TER-119), (all above antibodies were purchased from eBioscience). For human cell flow cytometry analysis, the following anti-human antibodies were used: CD31 (BD Pharmingen or eBioscience, clone WM59), CD34 (BioLegend, clone 561), CD45 (eBioscience or BioLegend, clone 2D1. BD, clone HI30), CD235a (BioLegend, clone HI264), and CD201 (EPCR, PROCR) (Miltenyi, clone REA337). Cell analysis and sorting were performed on LSR4, LSRII, FACSCantoII, FACSAria, and SORPAria flow cytometers (BD Biosciences). FlowJo software (BD Pharmingen) was used to analyze flow cytometry data.

For human quantitative PCR (qPCR) (Figure 4D), methylcellulose assay (MCA) (Figure S4C), and EC colony forming assay (Figure S4D), magnetically CD34 enriched fresh CBMNCs (by CD34 MicroBead Kit, Ultrapure, human, 130-100-453, Miltenyii, according to the manufacture's protocol) were incubated with anti-CD201 (EPCR, PROCR) (clone REA337) and anti-CD34 antibody (BioLegend, clone 561), sorted into 4 fractions; (1) CD34 High PROCR Low, (2) CD34 High PROCR High, (3) CD34 Bright PROCR Low, (4) CD34bright and PROCR high, performed on FACS Aria, and used for each experiment. This cell sorting was performed by FACS Aria and FACS Diva software (BD) was used for the sorting. The flow rate was set at 1.0.

For murine single EC culture, cells were sorted and each cell was directly loaded into each well of 96-well plate coated with OP9 stromal cell monolayer (see below). For endothelial cell colony forming limiting dilution assay, 20, 50, 100 P6 heart TdTomato$^+$ endothelial cells, or 200, 500, and 1,000 P6 lung TdTomato$^+$ endothelial cells were sorted into individual wells of OP9-coated 96-well plates, respectively.

**Endothelial colony formation culture**

For murine EC culture, OP9 stromal cells were maintained in OP9 medium (alpha-MEM medium (Gibco), with 20% FBS (Hyclone), and 0.5 % penicillin/streptomycin (Gibco)). To culture endothelial colonies, isolated murine endothelial cells were re-suspended in EC culture medium (alpha-MEM with 10% FBS, $5 \times 10^{-5}$ M β-mercaptoethanol (Sigma) and 0.5% penicillin/streptomycin (Gibco)). After 24 hours, non-adherent cells were removed by changing fresh medium. Medium was changed every 3 days afterwards until use. For human EC culture, the cells were re-suspended in complete EGM2 medium (Endothelial Basal Medium -2 (EBM-2, Lonza) with 10% FBS (Hyclone)) and re-plated on 0.1% type 1 rat tail collagen (Corning) coated tissue culture plates.

**Methylcellulose colony assay**

Methylcellulose colony assay was performed using MethoCult (H4434, StemCell Technologies) according to the manufacturer's protocol.

**3D *in vitro* tube forming assay in collagen gel**

200Pa stiffness pig skin type I collagen gels were made according to the manufacturer's instructions (Geniphys, Standarized Oligomer Polymerization Kit). Human platelet lysate (10%) (Sexton Biotechnologies) was then added into the gel and the liquid gel was kept on ice. Human EC were re-suspended in the gel at a density of ~$1 \times 10^6$ cells/1ml gel and each 50 μl gel was transferred to a well in 96-well plate. The cellularized gels were incubated at 37°C for 30 minutes to solidify. Next the gels were covered by adding 100 μl complete EGM2 medium to the wells. The cultures were checked under a microscope for every 12 hours until lumenized vessel-like structures were formed and identified.

***In vivo* gel implantation**

Blood from tamoxifen injected Tie2cre; ROSATdTomato pups were culture on OP9 for 3-

8 weeks. Then the cultured EC were collected and minced with a razor blade before mounting into collagen gels. Culture cells from individual pups were re-suspended in 250 μl 200pa collagen gel (Geniphys, Standarized Oligomer Polymerization Kit) plus 10% human platelet lysate (Cook) on ice and transferred into 1 well of 48-well plate. The gel was placed in a 37°C incubator to polymerize for 30 minutes. Next the cellularized gels were transplanted subcutaneously into the flanks of 6-12 weeks old NOD/SCID mice. The gels were retrieved from the animals at various time points between 14 days and 10 months post-implantation. To test the vessel forming potential of un-cultured mouse or human circulating ECFC, blood from 10 P3 Tie2Cre; ROSATdTomato pups (tamoxifen injected on P0, P1, P2), or MACS isolated 300,000 human cord blood $CD34^+$ cells were each suspended into a single collagen gel.

**Surgeries**

For EC collagen gel transplantation assay, cells were re-suspended in 250 μl 200pa pig skin type I collagen gel (Geniphys, Standarized Oligomer Polymerization Kit) plus 10% human platelet lysate on ice. When murine EC were tested, 50 μg/ml murine vascular endothelial growth factor (VEGF; Peprotech) and 100 μg/ml murine fibroblast growth factor 2 (FGF2; Peprotech) were added to the gels. Each cellularized gel was transferred into 1 well in a 48-well plate and incubated at 37 ˚C to polymerize for 30 minutes. Next the cellularized gels were transplanted subcutaneously into the flanks of 6-12 weeks old NOD/SCID mice as previously described (Prasain et al., 2014). The gels were retrieved from the animals at various time points between 14 days and 4 months post-implant.

**Cell culture immunohistochemistry and immunofluorescent staining**

For immunohistochemistry staining of endothelial colonies on OP9 co-culture plates, the cultures were fixed with 4% paraformaldehyde (PFA) for 30 minutes at RT. After washing, the samples were blocked with 2% skim milk (Sigma) in 0.1% triton (Sigma) PBS solution (PBSMT solution) for 30 minutes at RT and then stained with 1:100 rat anti mouse CD31 (BD Pharmingen, clone MEC 13.3) or rat anti mouse Flk1 (BD Pharmingen, clone Avas 12α1) antibody in PBSMT at RT for 2 hours or at 4°C overnight. Next the plates were stained with 1:200 alkaline phosphatase conjugated donkey anti rat IgG secondary antibody (Jackson ImmunoResearch) in PBSMT at RT for 2 hours or at 4°C overnight. The colonies were visualized by VECTOR Blue Alkaline Phosphatase (Blue AP) Substrate Kit.

For culture immunofluorescent staining of murine EC culture, fixed cultures were blocked with 10% goat serum in 0.5% triton PBS solution (blocking solution) and sequentially stained with primary antibody (1:100 rat anti mouse CD31, clone MEC 13.3) and secondary antibody (1:200 Alexa Fluor 488 or 647 conjugated goat anti rat IgG (Cell Signaling Technology)) in blocking buffer. All culture pictures were visualized using LeicaTM DM IL microscope with a SPOT RT3 camera (Spot Imaging).

**Tissue immunofluorescent staining**

To visualize Tdtomato$^+$ vessels in freshly collected muscle or collagen gel samples after transplantation, a Leica mz9.5 stereomicroscope with LEJ eqb 100 isolated lamp power supply was used. To detect the perfusion of implanted vasculature that had inosculated with host vessels, 100ul fluorescein conjugated Isolectin B4 (Vector Laboratories, for mice vessels) or 100ul fluorescein labeled Ulex Europaeus Agglutinin I (UEA I, Vector Laboratories, for human vessels) were intravascular injected into the mice 30 minutes prior to euthanization and sample collection. To take confocal images of tissues or transplanted gels, the samples were collected and fixed in 4% PFA at 4°C overnight, rinsed in 30% sucrose o at 4°C overnight, and then mounted in O.C.T. compound (Fisher Scientific) on dry ice. The tissue blocks were cut into 10-30 μm sections using a Leica CM3050s cryostat and mounted on Superfrost Plus Gold microscope glass slides (Thermo Fisher Scientific). After blocking with blocking buffer at RT for 1 hour, the slides were then stained with different unconjugated primary anti bodies include: rat anti mouse CD31 (BD Pharmingen, clone MEC 13.3, 1:100), and rabbit anti ERG (Abcam, clone EPR3864. 1:100), at 4°C overnight. Then 1:200 Alexa Fluor 488, 555, or 647 conjugated goat anti-rat, anti-rabbit, or anti-mouse IgG antibodies (Cell Signaling Technology) were used for secondary staining at 4°C overnight. For some stainings, the following conjugated antibodies were used: Alexa Fluor 647 conjugated mouse anti human CD31 (BD Pharmingen, Clone WM59. 1:50), Alexa Fluor 488 or 594 conjugated mouse anti smooth muscle actin α (eBioscience, clone 1A4. 1:100). After staining, the samples were mounted with ProLong Gold Antifade Mountant with DAPI (Molecular Probes) and Z-stack confocal images were taken on Olympus FV1000 microscope.

All fluorescent pictures were processed using ImageJ software to produce merge images. 3D reconstruction of $CD31^+$ and $TdTomato^+$ vessels in tissues or gels were performed using Imaris software. The volumes of blood vessels were calculated by Imaris software using "Surface" function according to the manufacturer's instruction.

**Quantitative PCR**

RNA from each sample was extracted using RNeasy Plus Micro kit (Qiagen). Reverse transcription was done using SuperScript IV Reverse Transcriptase (ThermoFisher

Scientific). Quantitative PCR was performed on StepOnePlus (ThermoFisher Scientific) with PowerUp SYBR Green Master Mix (Applied Biosystems). Beta-actin was used as reference gene to calculate transcript abundance of each target gene. The expression level fold change between sample genes and reference genes were calculated by standard curve, where for every run, a new standard curve was constructed.

### Statistical analyses

Unless otherwise mentioned, all data are presented as mean ± standard deviation and unpaired two tailed Student's t-test was used to determine significance. If p value > 0.05, differences considered non-significant (n.s.), while significance was confirmed and marked: *: p < 0.05, **: p < 0.01, ***: p < 0.001, or ****: p < 0.0001. All statistical analysis were calculated using Graphpad Prism or Microsoft Excel software.

Aibar, S., Gonzalez-Blas, C.B., Moerman, T., Huynh-Thu, V.A., Imrichova, H., Hulselmans, G., Rambow, F., Marine, J.C., Geurts, P., Aerts, J., et al. (2017). SCENIC: single-cell regulatory network inference and clustering. Nat Methods *14*, 1083-1086. 10.1038/nmeth.4463.

Benz, C., Martins, V.C., Radtke, F., and Bleul, C.C. (2008). The stream of precursors that colonizes the thymus proceeds selectively through the early T lineage precursor stage of T cell development. J Exp Med *205*, 1187-1199. 10.1084/jem.20072168.

Dobin, A., Davis, C.A., Schlesinger, F., Drenkow, J., Zaleski, C., Jha, S., Batut, P., Chaisson, M., and Gingeras, T.R. (2013). STAR: ultrafast universal RNA-seq aligner. Bioinformatics *29*, 15-21. 10.1093/bioinformatics/bts635.

Epelman, S., Lavine, K.J., Beaudin, A.E., Sojka, D.K., Carrero, J.A., Calderon, B., Brija, T., Gautier, E.L., Ivanov, S., Satpathy, A.T., et al. (2014). Embryonic and adult-derived resident cardiac macrophages are maintained through distinct mechanisms at steady state and during inflammation. Immunity *40*, 91-104. 10.1016/j.immuni.2013.11.019.

Hochstetler, C.L., Feng, Y., Sacma, M., Davis, A.K., Rao, M., Kuan, C.Y., You, L.R., Geiger, H., and Zheng, Y. (2019). KRas(G12D) expression in the bone marrow vascular niche affects hematopoiesis with inflammatory signals. Exp Hematol *79*, 3-15 e14. 10.1016/j.exphem.2019.10.003.

Madisen, L., Zwingman, T.A., Sunkin, S.M., Oh, S.W., Zariwala, H.A., Gu, H., Ng, L.L., Palmiter, R.D., Hawrylycz, M.J., Jones, A.R., et al. (2010). A robust and high-throughput Cre reporting and characterization system for the whole mouse brain. Nat Neurosci *13*, 133-140. 10.1038/nn.2467.

Muzumdar, M.D., Tasic, B., Miyamichi, K., Li, L., and Luo, L. (2007). A global double-fluorescent Cre reporter mouse. Genesis *45*, 593-605. 10.1002/dvg.20335.

Novershtern, N., Subramanian, A., Lawton, L.N., Mak, R.H., Haining, W.N., McConkey, M.E., Habib, N., Yosef, N., Chang, C.Y., Shay, T., et al. (2011). Densely interconnected transcriptional circuits control cell states in human hematopoiesis. Cell *144*, 296-309. 10.1016/j.cell.2011.01.004.

Prasain, N., Lee, M.R., Vemula, S., Meador, J.L., Yoshimoto, M., Ferkowicz, M.J., Fett, A., Gupta, M., Rapp, B.M., Saadatzadeh, M.R., et al. (2014). Differentiation of human pluripotent stem cells to cells similar to cord-blood endothelial colony-forming cells. Nat Biotechnol *32*, 1151-1157. 10.1038/nbt.3048.

Stuart, T., Butler, A., Hoffman, P., Hafemeister, C., Papalexi, E., Mauck, W.M., 3rd, Hao, Y., Stoeckius, M., Smibert, P., and Satija, R. (2019). Comprehensive Integration of Single-Cell Data. Cell *177*, 1888-1902 e1821. 10.1016/j.cell.2019.05.031.

Trapnell, C., Cacchiarelli, D., Grimsby, J., Pokharel, P., Li, S., Morse, M., Lennon, N.J., Livak,

K.J., Mikkelsen, T.S., and Rinn, J.L. (2014). The dynamics and regulators of cell fate decisions are revealed by pseudotemporal ordering of single cells. Nat Biotechnol *32*, 381-386. 10.1038/nbt.2859.

Zhao, Y., Li, X., Zhao, W., Wang, J., Yu, J., Wan, Z., Gao, K., Yi, G., Wang, X., Fan, B., et al. (2019). Single-cell transcriptomic landscape of nucleated cells in umbilical cord blood. Gigascience *8*. 10.1093/gigascience/giz047.
